# Structurally diverse calloses/β-1,3-glucans in plant cell wall microdomains

**DOI:** 10.1101/2024.07.04.602027

**Authors:** Sam Amsbury, Susan E. Marcus, Richa Yeshvekar, Jenny Barber, Liam German, James F. Ross, Ieva Lelenaite, Tatiana de Souza Moraes, Janithri Wickramanayake, Anastasiya Klebanovych, Kirk Czymmek, Tessa M. Burch-Smith, Emmanuelle M. Bayer, William Willats, Iain W. Manfield, Paul Knox, Yoselin Benitez-Alfonso

**Affiliations:** Plants, Photosynthesis and Soil, School of Biosciences, The University of Sheffield; Sheffield, United Kingdom; Centre for Plant Sciences, School of Biology, University of Leeds; Leeds LS2 9JT, United Kingdom; Astbury Centre for Structural Molecular Biology, University of Leeds; Leeds LS2 9JT, United Kingdom; School of Natural and Environmental Sciences, Newcastle University; Newcastle upon Tyne, United Kingdom; Laboratoire de Biogenèse Membranaire, UMR5200, CNRS, Université de Bordeaux; Villenave d’Ornon, France; Advanced Bioimaging Laboratory, Donald Danforth Plant Science Center; St. Louis, Missouri, 63132, USA; Donald Danforth Plant Science Center; St. Louis, Missouri, 63132, USA

**Author notes:** Crop Science Centre, Department of Plant Sciences, University of Cambridge; 93 Lawrence Weaver Road, Cambridge CB3 0LE, United Kingdom (Richa Yeshvekar, Liam German).

**Keywords:** callose, β-(1,3)-glucan, glycan polymers, monoclonal antibodies, cell walls, polysaccharides, plants

## Abstract

Cell walls underpin the mechanics of cell growth, intercellular signalling, and defence against pathogenic organisms. β-(1,3)-glucans (also known as callose) are polysaccharides found in plants, fungi, and some bacterial species. In developing plant organs, callose accumulates around intercellular channels (plasmodesmata) controlling cell-to-cell communication. We developed monoclonal antibodies for the detection of β-(1,3)-glucans and using these identified distinct populations of callose differing in size and secondary structure. Callose sub-populations were in proximal but not overlapping cell wall microdomains implying distinct spatial and functional microenvironments. We also unveiled callose interaction with xyloglucan; another plant glycan regulating cell wall functions. This work challenges previous views demonstrating structural heterogeneity in plant callose and supporting interactions between glycans with roles in the regulation of cell wall properties and functions.

---

Cell walls are crucial for organismal development and defence (*1-4*). They consist of a diverse array of polysaccharides that interact and assemble into dynamic glycan networks. In plant cell walls, the glycan network is formed primarily by cellulose microfibrils, pectins, and other matrix polysaccharides (including xyloglucans, xylans and mannans). Understanding the structures of these polysaccharides is essential for deciphering their roles in cell wall mechanics and development. Past research has revealed the complex interactions between major cell wall components, where modifications such as pectic homogalacturonan methyl esterification can significantly impact cellulose microfibril organization, cell wall physical properties and function (*5-7*). These insights highlight cell walls as a key target for improving crop growth and resilience to climate change.

β-(1,3)-glucans (known as callose) are a minor component in plant primary cell walls with their structural properties remaining poorly understood (*4, 8-10*). While major β-(1,3)-glucan sources reside in fungal and bacterial cell walls (*11-13*), in plants, callose accumulates in specialized cells and interfaces including pollen tube walls (*14*), cell plates (*2*), sieve pores (*15*) and surrounding membranous intercellular channels named plasmodesmata (*4, 10, 16-18*). Despite its low abundance, callose plays diverse and critical roles in plant development: forming protective papillae during pathogen attacks, controlling pollen development and thereby fertilization, and regulating cell-to-cell signalling and the transport of developmental proteins, RNAs and hormones via plasmodesmata (*2, 4, 9, 14, 19, 20*). However, our knowledge of callose’s physical properties, interactions with other cell wall components, and its overall impact on cell wall functions is limited (*8, 10*).

Here we dissect the biochemical/structural properties and interactions of callose. We report on the identification of subpopulations of callose that can be distinguished by size and charge using new monoclonal antibodies (mAbs). Structurally heterogenous calloses are regulated by callose synthases, bind distinct cell wall microdomains at plasmodesmata and interact differently with xyloglucan suggesting a role in regulating cell wall properties and function.

## Results

### Monoclonal antibodies identify structurally distinct calloses

Callose detection relies on aniline blue staining and immunofluorescence with a commercial antibody (Biosupplies, here named BioS) (*21, 22*) but the size, branching and/or secondary structure of the polymer are unknown.

To dissect callose structures, we carried out a biochemical separation of Arabidopsis cell walls. Alcohol-insoluble residues were sequentially extracted to release weakly (soluble in trans-1,2-Diaminocyclohexane-N,N,N’,N’-tetraacetic acid hydrate, CDTA) and strongly (solubilized by potassium hydroxide, KOH) bound cell wall polysaccharides (*23*). The KOH extract was selected for further fractionation using Size Exclusion Chromatography (SEC) (Figure 1A). Each fraction was probed with BioS in an Enzyme-Linked Immunosorbent Assay (ELISA) to obtain the SEC profile. Two major populations of calloses were observed, eluting at retention volumes corresponding with a 40 kDa and a 20 kDa molecular weight dextran standards (Fig. S1).

**Fig. 1.**
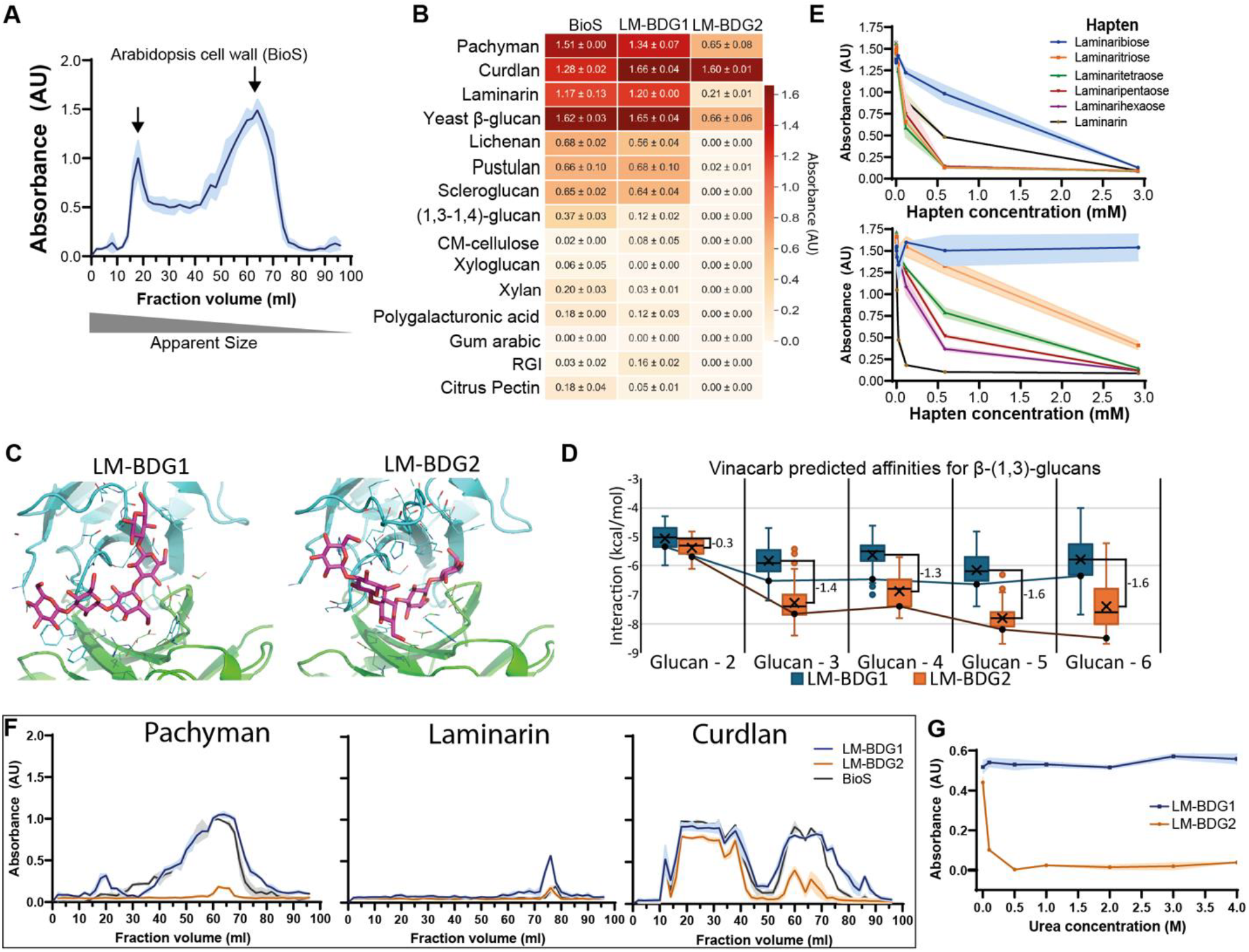
Monoclonal antibodies recognize distinct β-(1,3)-glucan epitopes. (**A**) Size exclusion chromatography fractions of Arabidopsis cell walls (KOH-extracted fraction) were probed using ELISA and anti-β-(1,3)-glucan from Biosupplies (Bios). ELISA absorbance traces show mean ± SD (shading), n=2 pools of 8 plants. Arrows indicate two main peaks of elution representing callose subpopulations. (**B**) Novel rat monoclonal antibodies (mAbs LM-BDG1 and LM-BDG2), used in glycomic analysis alongside BioS, display no significant cross reactivity with non-β-(1,3)-glucans (in red box). LM-BDG1 and BioS have strong affinity for all major β-(1,3)-glucans whilst LM-BDG2 shows strong binding to curdlan and weak binding to pachyman, laminarin and yeast β-glucan. Data show mean ± SD, n=5. RGI = rhamnogalacturonan-I. (**C)** LM-BDG1 and LM-BDG2 predicted structures (blue and green, AlphaFold2) after docking with a β-(1,3)-glucan pentasaccharide (magenta, low energy conformation) docked with Vina-Carb (ligand extracted from PDB_2W62). (**D**) Vina-Carb predicted binding affinities in relation to glucan sizes. Low energy cluster centres are highlighted with black circle. (**E**) Hapten inhibition assays for LM-BDG1 and LM-BDG2 using increasing concentrations of oligo-β-(1,3)-gluco-saccharides from a disaccharide (laminaribiose) to a hexosaccharide (laminarihexaose) and alongside laminarin. (**F**) Binding of mAbs to β-1,3-glucan standards: pachyman, laminarin and curdlan. LM-BDG1 reveals multiple populations in both pachyman and curdlan. LM-BDG2 binds predominantly to curdlan, with a strong preference for the large sized polymers. (**G**) LM-BDG1 and LM-BDG2 detection of curdlan pre-treated with different concentrations of urea prior to coating ELISA microtitre plates. LM-BDG2 binding to curdlan is rapidly disrupted suggesting conformation dependence. Shaded portions indicate standard deviation from mean, n=3.

To characterise callose sub-populations, mAbs were generated using rat hybridoma technology (*24*). Two lines, named Leeds-Monoclonal-β-(1,3)-D-Glucan 1 and 2 (LM-BDG1 and LM-BDG2), were selected for detailed characterisation due to their differing binding to β-(1,3)-glucan variants in relation to BioS (Figure 1B and Figure S2). In ELISAs, both LM-BDG1 and BioS gave strong signals with pachyman, curdlan, laminarin and yeast β-glucan and moderate-weak signals with lichenan, pustulan and scleroglucan. All these substrates contain β-(1,3)-glucosyl residues and differing number of β-(1,6)-side chain (*11*). BioS (but not LM-BDG) also binds weakly β-(1,3-1,4)-glucan (Figure 1B). LM-BDG2 binding differed distinctly from LM-BDG1 and BioS, with highest signal obtained with curdlan, moderate signal with pachyman and yeast β-glucan and weak signal with laminarin (Figure 1B). LM-BDG2 did not display any cross-reactivity to non-β-(1,3)-glucan polymers (i.e., CM-cellulose, xyloglucan, xylan, rhamnogalacturonan (RG) I, or citrus pectin). LM-BDG1 and BioS, on the other hand, bind very weakly some of these (Figure 1B). In general, LM-BDG mAbs display a higher specificity than Bios in ELISA assays.

To confirm our results, glycomicroarray analyses were carried out (*25, 26*) (Figure S2). Although there were differences in relative avidities, all three mAbs were highly specific for β-(1,3)-glucan polymers with very little or no binding to an extended set of control polysaccharides including pectins, gums, and other non-β-(1,3)-glucan (Figure S2). In contrast to ELISA, LM-BDG2 binding to carboxymethyl-curdlan was slightly weaker than for BioS and LM-BDG1 (Figure S2). The glycomicroarray data also revealed a relatively strong binding of LM-BDG1 for barley and oat β-glucans. An additional mAb, named LM-BDG3, was also characterised using glycomicroarrays (Figure S2). LM-BDG3 had a very similar binding profile to LM-BDG1, but with reduced binding to barley and oat β-glucans.

To further dissect structural differences, we sequenced the variable regions of LM-BDG1 and LM-BDG2 and computationally predicted their structure and docked glucans of different sizes (*27*). The structures were predicted by AlphaFold (*28*), yielding a global per residue Local Distance Difference Test (pLDDT) score of 94.4 for LM-BDG1 and 95.4 for LM-BDG2, both in the ‘very high confidence’. A small loop (β-hairpin) in the binding cleft for each model fell to a pLDDT of ∼80, however, given the limited conformational options for β-hairpins, and the high overall pLDDT scores locally and globally, these structures were deemed of adequate quality to perform ligand docking [see methods]. Vina-Carb (*29*) generated an ensemble of docked poses for a range of 2 to 6 residue β-1-3-gluco-oligosaccharides (glucan). For each protein with each oligosaccharide, an interface energy landscape was computed based on Vina-Carb affinities superimposed on to clustered ligand RMSD relationships (root mean square deviation). The results highlight the occupancy and average binding affinity per cluster grouping (Figure S3). An example conformation predicted for glucan-5 is shown in Fig.1C. Plotting the interaction energies of the different binding conformations revealed that while LM-BDG1 values remain largely unchanged across the different ligand sizes, LM-BDG2 binding energy is reduced as glucan size increases suggesting a higher affinity for larger ligands (Fig. 1D).

Hapten inhibition assays, where binding to an immobilised β-(1,3)-glucan (i.e., curdlan) is inhibited by increasing concentrations of β-(1,3)-gluco-oligosaccharides, were carried out to define experimentally minimal epitope recognition (Figure 1E). LM-BDG1 binding suggests minimal epitope recognition of a β-(1,3)-trisaccharide (laminaritriose). Oligosaccharides weakly inhibited LM-BDG2 binding which was only fully inhibited with laminarin (of ∼25 glucoses (*30*)) (Figure 1E). The results confirm LM-BDG2’s relative higher affinity for large glucans.

The size dependence was further investigated using SEC separation of commercially available β-(1,3)-glucan standards (Fig. 1F). Pachyman from the fungi *Poria cocos* has a degree of polymerization (DP) of ∼255, curdlan from *Alcaligenes faecalis* has a DP that ranges between 155 - 450, whereas laminarin (DP ∼25) is extracted from *Laminaria digitata* (*12, 30, 31*). SEC profile for LM-BDG1 and BioS was very similar detecting elution peaks for pachyman, laminarin and curdlan. Curdlan eluted in two main populations but LM-BDG2 preferentially bound the larger size fraction (10-30 ml elution volume) supporting its higher affinity for larger glucans (Fig. 1F).

Curdlan forms single and triple helical structures due to intramolecular and intermolecular hydrogen bonding (*12, 31*). To test if LM-BDG affinities for curdlan is related to its complex structural conformations, we used urea which disrupt glycan structures (*32*) (Figure 1G). ELISAs indicated a loss in LM-BDG2 binding to curdlan after treatment with 0.5 M urea whereas LM-BDG1 binding is lost after treatment with a 4 M concentration.

The results indicate structural differences in callose structures bound by LM-BDG1 and LM-BDG2 underpinning the differing epitope specificities.

### Structurally heterogenous callose revealed at plasmodesmata cell wall microdomains

Commercial β-(1,3)-glucans display drastically differing physical properties, such as solubility and gelling properties underpinning their mechanical and functional behaviour (*11, 12, 30, 31*). We propose structural diversity in calloses as an important aspect of its functions. To validate the existence of structurally heterogeneous calloses *in planta*, we performed fluorescence immunolocalization on a range of plant materials (Figure 2 and Figure S4, S5). LM-BDG1 and LM-BDG2 specifically targeted distinct but overlapping subcellular locations: pit fields in tomato fruit pericarp (Figure 2A-D, Fig. S4), Arabidopsis guard cells (Fig. 2E-L) and cell walls associated with plasmodesmata in aspen buds and tobacco root cells (Figure S5). These binding patterns coincide with regions previously reported to contain callose (*10, 17, 21, 33*). To confirm their affinity for plasmodesmatal callose, we tested Arabidopsis leaves enriched in callose by ectopic expression of the Plasmodesmata-Callose Binding protein 1 (PDCB1-OE, (*34*)) (Fig. 2M-N). Both LM-BDG1 and LM-BDG2 bound to large aggregates in PDCB1 overexpressing leaves not seen in wildtype controls (Fig. S4). In all cases, LM-BDG2 displayed a more restricted distribution, suggesting recognition of a less abundant epitope. To link the punctate binding pattern with plasmodesmata locations, we carried out immunogold-transmission electron microscopy in Arabidopsis cell cultures, aspen meristems (Fig. S6) and in tobacco leaf sections (Fig. 3A-B and Fig. S7). The LM-BDG2 epitope was detected with low abundance at some (not all) plasmodesmata locations in tobacco leaf and aspen meristem sections. Labelling with this antibody was absent in Arabidopsis cell cultures (Fig. S6). LM-BDG1 gold particles, on the other hand, decorated most cell walls at the plasmodesmatal region in all studied plant species.

**Fig. 2.**
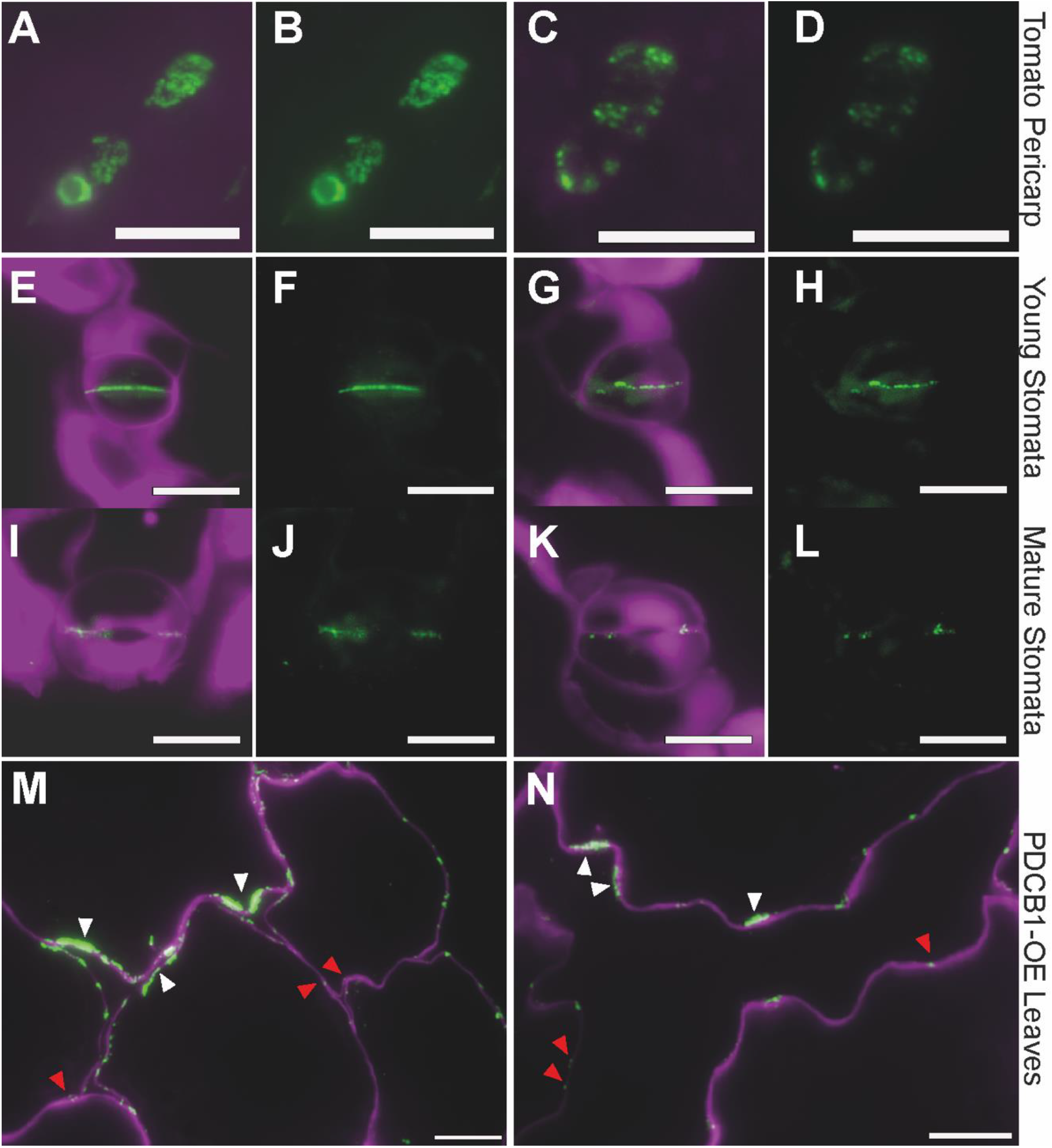
Immunofluorescence labelling of callose populations in plant material revealed differing profiles. LM-BDG1 (**A-B, E-F, I-J, M)** and LM-BDG2 (**C-D, G-H, K-L, N)** labelled callose in pit fields from tomato fruit pericarp (**A-D**), in Arabidopsis young (**E-H**) and mature (**I-L**) leaves stomata, and in leaf sections of the PDCB1 overexpression (PDCB1-OE) line – which is known to accumulate callose (**M-N**, control are included in Fig. S4). Antibody binding was revealed using Alexa-488 anti-rat secondary antibody (green signal). Cell walls were counterstained with calcofluor white (shown in magenta) in merged panels **A, C, E, G, I, K, M-N**. Punctate pattern is indicative of plasmodesmata (red arrows). PDCB1-OE also accumulates callose in large aggregations, indicative of ectopic expression (white arrows). Scale bars = 10 µm.

**Figure 3.**
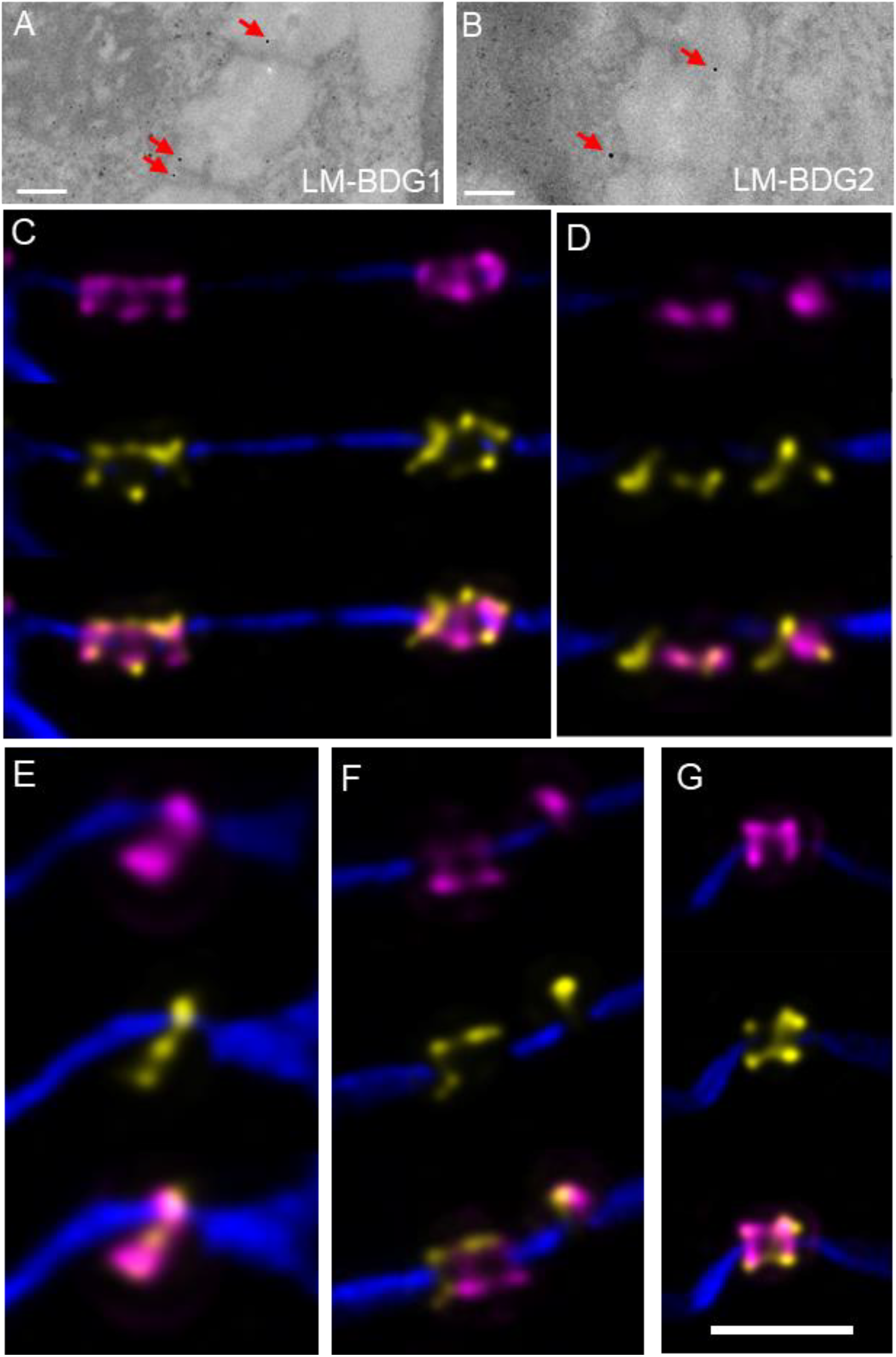
High resolution sequential multiplex labelling of callose reveals distinct and partially overlapping binding microdomains. (**A-B**) Immunogold EM labelling in tobacco leaf sections shows that callose (red arrows) is detected by both LM-BDG1 and LM-BDG2 at plasmodesmata (gold particles appear around the darker electron dense regions crossing cell walls). Scale bar = 200 nm. (**C-G**) Immunofluorescence localization reveals binding of LM-BDG1 (in magenta) and LM-BDG2 (in yellow) at cell wall microdomains presumed plasmodesmata. Calcofluor counterstain of cell walls is shown in blue. The bottom picture in each panel shows the merged channels. LM-BDG1 and LM-BDG2 epitopes display strong overlap but do not occupy the exact same space. (**E-G**) The epitopes display a range of distributions at plasmodesmata. LM-BDG1 can accumulate at either side of a presumed single plasmodesmata pore (**E**) while LM-BDG2 is distributed along the length of the pore. In complex/twinned plasmodesmata (**F**,**G**) both epitopes are similar in abundance, sometimes distributed asymmetrically, on either side of the pores. Scale bar (**C-G**) = 300 nm.

For a more detailed visualisation of the relative epitope distributions, we carried out lattice super-resolution (SIM^2^) and multiplex microscopy in tobacco leaf sections (Fig.3 and S7). We labelled with LM-BDG1, stripped, and subsequently label with LM-BDG2 (and in reverse order). Although both monoclonal antibodies broadly labelled leaf plasmodesmata, the fluorescence signals did not completely overlap, indicating that they bind to closely related but separate cell wall microdomains (Fig. 3C-G). The LM-BDG1 epitope was more regularly distributed at plasmodesmata, including locations where the LM-BDG2 epitope was absent (Fig S7). Both epitopes were detected in what looked like simple and branched plasmodesmata (Fig 3C-D) with LM-BDG1 typically binding a larger (and neck) region compared to LM-BDG2.

LM-BDG2 labelling can occur along the entire channel length (Fig 3E) and sometimes appear distributed differently on opposing sides of the cell walls (Fig 3F,G).

Collectively, these findings demonstrate the presence of structurally different callose epitopes with varying abundances in cell walls from multiple plant species.

### Differential regulation of calloses in cell wall extracts and by callose synthase

The structural differences detected by LM-BDG1 and LM-BDG2 may point to callose structures that can interact and be regulated differently within cell walls. To dissect the properties and potential interactions of callose epitopes in cell walls, we fractionated Arabidopsis cell wall extracts using SEC and probed the fractions with LM-BDG1 and LM-BDG2 (Fig. 4). Sequential extraction of alcohol-insoluble residues with CDTA and KOH, was followed by enzymatic digestion to release cellulose-microfibril interacting polymers. CDTA extracts yielded a single, mid-sized callose population identified only by LM-BDG1. KOH extraction revealed two major callose subpopulations of relatively large and small size (∼ 20 ml and ∼65 ml retention volumes, see Fig.1) when probed with LM-BDG1 and BioS. As expected, LM-BDG2 specifically recognised epitopes present in the larger subpopulation. Some mAb signal, not resolved into clear peaks, was detected between these two main peaks indicating a small amount of intermediate sized callose. Cellulase treatment released callose populations of various sizes, with peaks at various retention volumes. Notably, in the cellulase extract LM-BDG2 bound the smaller populations (∼60 and 80ml) indicating that polysaccharide size alone does not govern epitope recognition and, more likely, this depended on structural conformation (Fig. 4). The small size of the LM-BDG2 bound population in the cellulase extract may represent unstructured chains of callose interacting with cellulose microfibrils and released after digestion.

**Fig. 4.**
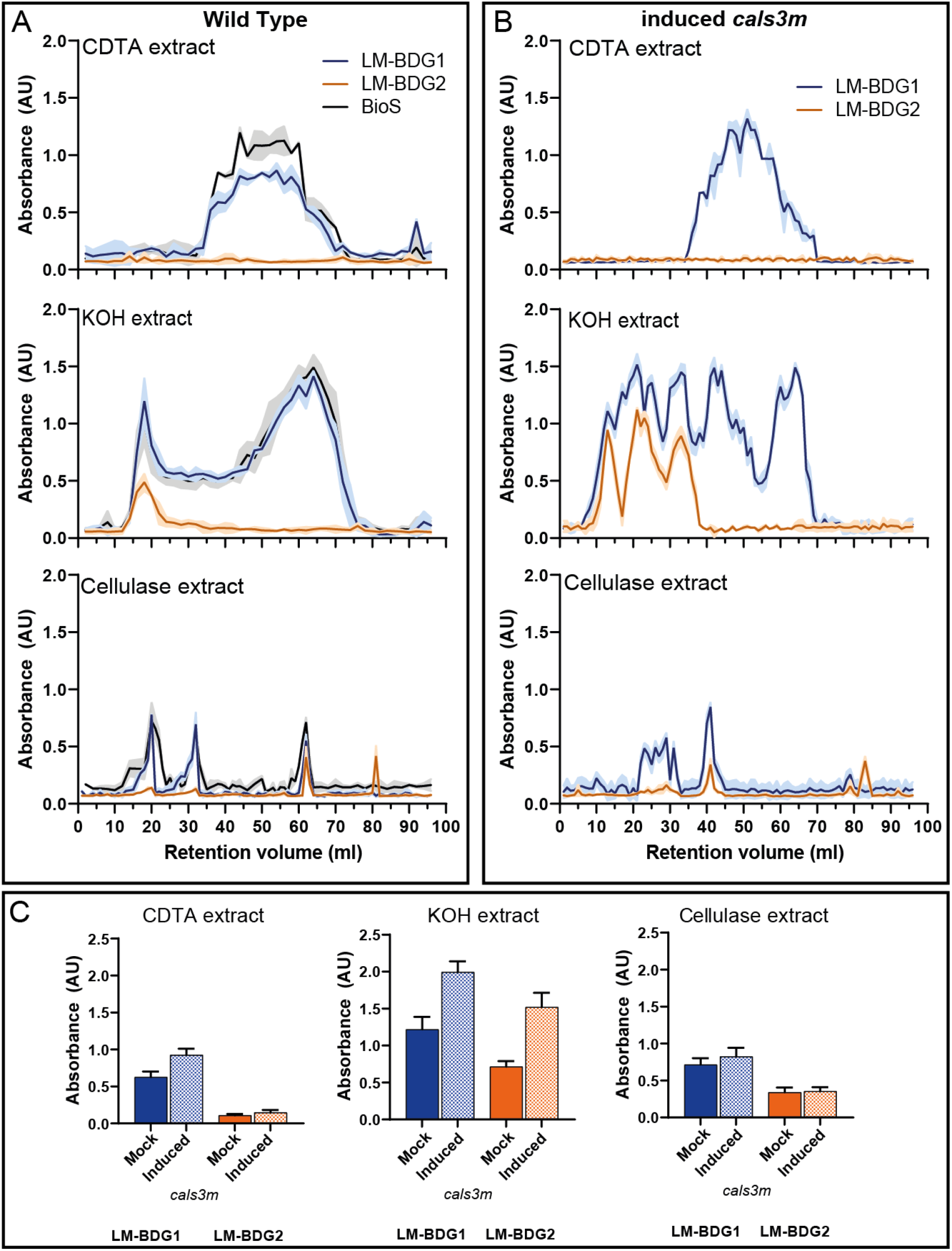
Chromatographic separation and antibody detection profiles indicate diverse populations of callose in plant cell wall extracts. Size exclusion chromatography was used to separate *Arabidopsis thaliana* cell walls sequentially extracted with CDTA, KOH and the remaining pellet digested with cellulase (cellulase extract). Profiles are absorbance traces after ELISA-based immunodetection with LM-BDG1 (blue trace), LM-BDG2 (orange trace) and the BioS (black trace). Chromatographs show mean ± SD (shading), n=2 pools of 6 plants. (**A**) CDTA-extracted callose shows a single mid-sized population that only binds LM-BDG1 and BioS. KOH extraction reveals two callose subpopulations: large (eluting at ∼20 ml) detected by LM-BDG1, BioS and LM-BDG2 and small (eluting at ∼65 ml) detected only by LM-BDG1 and BioS. Cellulase treatment releases callose of varying sizes detected by all antibodies with LM-BDG2 binding only the small callose population (eluting at 60 and 80 ml). (**B**) Induction of callose synthesis by ectopic expression of a mutant callose synthase (*cals3m*) increased the levels of callose in the CDTA and KOH extracts and altered size profiles in both KOH and cellulase-digested fractions. LM-BDG2 binding was not detected in the CDTA fraction. **(C)** ELISA data showing total callose content in mock and induced *calsm3* plants. Note signals are higher in induced *cals3m* (mainly in the KOH fraction) for both LM-BDG1 (blue bars) and LM-BDG2 (orange bars) antibodies. Error bars represent standard error of the mean (SEM) from n=4 pools of 4 plants each.

Ectopic induction of callose by expression of a hyperactive (mutated) version of *Arabidopsis thaliana* callose synthetic enzyme CALS3 (named *cals3m*) reduces plasmodesmata transport and alters cell wall composition (i.e., cellulose microfibril organization), hydrophilicity and digestibility (*17, 33, 35*). To investigate how CALS3 activity affects epitope abundance, we compared the SEC profiles from wildtype and induced *cals3m* cell walls. As expected, increased callose accumulation was detectable in induced *cals3m* by both LM-BDG1 and LM-BDG2 (Fig. 4C), along with changes in size distribution (Fig. 4B). As in wildtype, LM-BDG2 only bound large glucans fractionated with KOH and small sizes extracted in cellulase-digested fractions. In the KOH-extracted fractions, additional intermediate peaks indicated accumulation of a broader range of callose sizes. No significant increase in callose released after cellulase digestion was detected in *cals3m* plants but the size profile was substantially altered compared to wildtype.

The results from *cals3m* induction suggest that dynamic callose metabolism plays a role in modulating the abundance of different callose structures. The release of callose post-cellulose digestion supports callose-cellulose interactions *in muro*. Such interactions potentially influence, and are influenced by, callose structural heterogeneity with impacts on cell wall behaviours.

### Analysis of callose subpopulations reveals novel interactions with xyloglucan

In primary cell walls, pectin and other polysaccharides, such as xyloglucan, co-exist with cellulose and may play a role in regulating callose structures or functions. We carried out enhanced SEC separations, using a longer elution profile than before, to better resolve the callose peaks. We focused the analysis on the KOH fraction which extracted most of the callose polymers (Fig. 4). Besides LM-BDG1 and LM-BDG2, we probed the same fractions with LM19 (mAb directed to pectic homogalacturonan (*24*)) and LM25 (mAb directed to xyloglucan (*36*)) to dissect potential co-elution patterns, indicative of interacting polymer networks (*23*).

Enhanced SEC analysis (Fig. 5A) resolved peaks corresponding to a range of callose sizes co-eluting with xyloglucan (grey arrows), and with both xyloglucan and homogalacturonan (red arrows). While large size callose detected by both LM-BDG1 and LM-BDG2, co-eluted with both pectin and xyloglucan, the smaller size population (detected by LM-BDG1 alone) mainly co-eluted with the xyloglucan-containing fractions. These results suggest that callose might associate with xyloglucan and/or with pectin potentially forming pectin-xyloglucan-callose complexes of larger sizes. To dissect further, we applied anion-exchange chromatography (AEC) which separates fractions using a salt gradient (*23*). Pectins are negatively charged and interact with xyloglucans (a neutral polysaccharide) forming an anionic complex that elute at a higher volume in AEC profiles. Three peaks (two in the neutral and one in the acidic region) were detected after probing KOH extracts fractionated using AEC with LM-BDG1. A negatively charged peak (eluting at 60 ml) coincided with a broad peak detected by LM19 suggesting potential for callose-pectin interactions (Fig. 5B). The callose profile though displayed closer resemblance to the LM25 profile indicating co-elution of callose and xyloglucan in both the neutral population and in the charged population (Fig. 5B).

**Fig 5.**
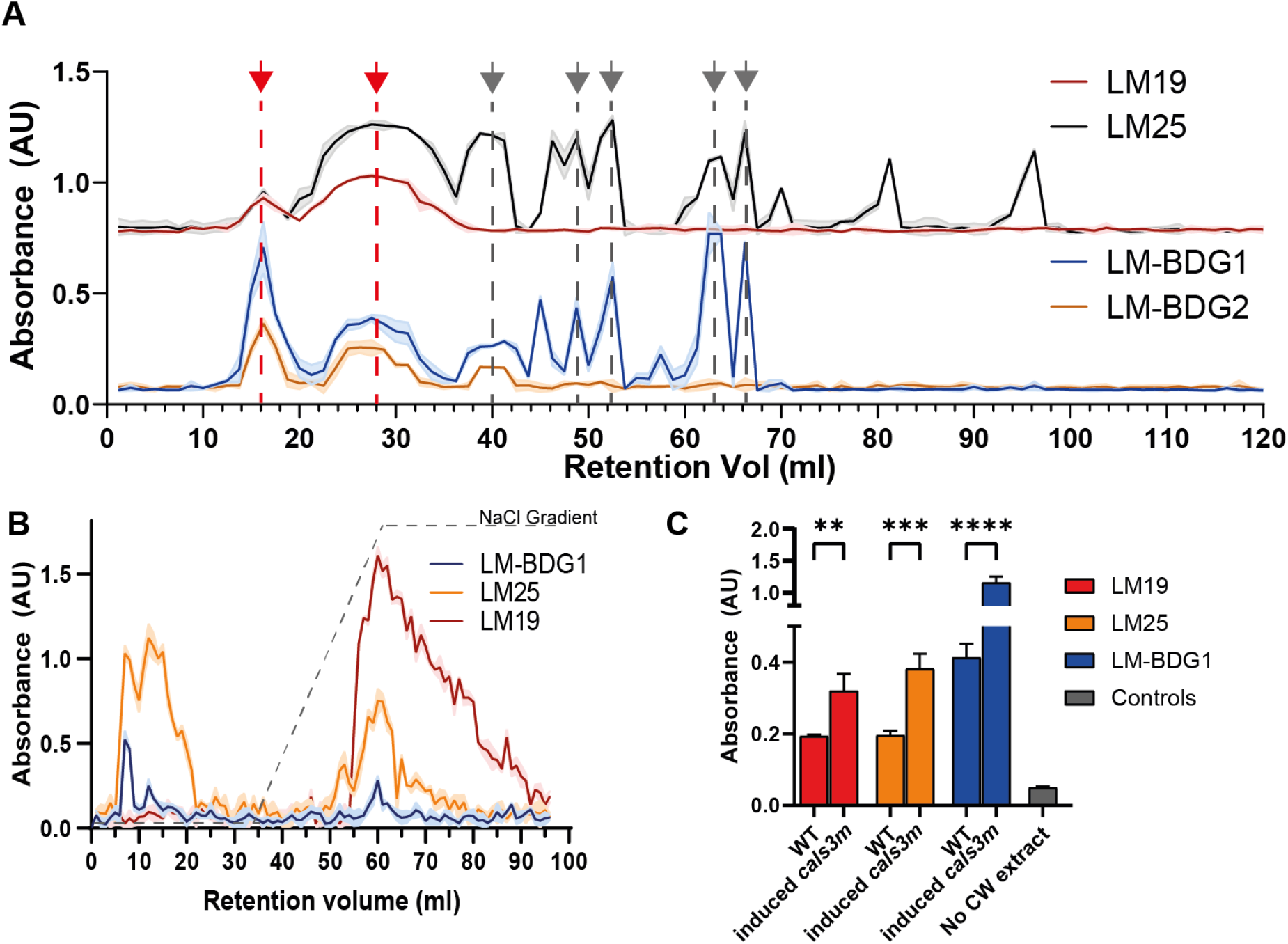
Interactions between callose and cell wall polysaccharides revealed by chromatographic separations and sandwich ELISAs. (**A**) Enhanced size-exclusion chromatography of KOH-extracted WT Arabidopsis cell walls revealed that LM-BDG1-bound callose co-elutes with xyloglucans (detected using the LM25 antibody, grey arrows) and/or pectins (detected using the LM19 antibody, red arrows). LM-BDG2 mainly binds calloses co-eluting with both xyloglucans and pectins (red arrows). (**B**) Anion exchange chromatography of KOH-extracted WT cell walls reveals elution of LM-BDG1-bound callose (blue trace) in two small neutral peaks (that co-elutes with xyloglucans/LM25) and in an acidic peak (that co-elutes with both xyloglucans and pectins/LM19) suggesting callose-xyloglucan-pectin interactions. (**C**) Sandwich ELISA experiments carried out using a callose-binding module (CBM43) to capture callose from KOH-extracts. Signal (higher than the no cell wall/ CW extract control) was detected when the CBM43-bound callose was probed with the LM19 (pectin) and LM25 (xyloglucan) antibodies indicating glycan interactions in both wildtype (WT) and in cell walls extracted from *cals3m* induced line. Error bars and shaded areas represent standard error of the mean (SEM) from n=4 pools of 4 plants each. Statistical significance: ** indicates p<0.05 *** indicates p<0.005 *** indicates p<0.0005 determined by ANOVA with Holm multiple comparisons correction.

Co-elution of epitopes in both SEC and the AEC profiles strongly suggest interactions between callose-xyloglucan and pectin polymers. We further tested this hypothesis using sandwich-ELISAs, where a first, immobilised mAb captures a target molecule, and a second mAb is used to detect components that bind with the targeted antigen (*37*). After washing, binding by the second mAb may only occur when polysaccharides interact through hydrogen bonding, close entanglements or covalent associations. We used a previously described callose-binding carbohydrate-binding module (CBM43) to capture callose (*34*) and then probed the bound material with LM25 and LM19 to detect xyloglucan and pectin respectively (Fig. 5C). CBM43 was able to bind callose (detected with LM-BDG1) from wildtype and induced *cals3m* cell walls. Probing the CBM43-bound fractions with both LM19 and LM25 revealed pectin and xyloglucan association with callose. Increasing callose amount in induced *cals3m* led to increase in LM19 and LM25 binding supporting these associations. To verify these results, we reversed the analyses using LM25, LM19 and a different callose antibody (LM-BDG3, Fig.S2) to capture respective antigens and a purified HRP-tagged version of LM-BDG1 to detect calloses. Callose was mainly detected, in both the KOH and the cellulase extracts, with the LM25-bound antigen and only weakly in the LM19-bound antigen (Fig. S8 B-E). Together, the results are indicative of strong interactions between callose and xyloglucan, potentially alone and in complexes with pectin.

## Discussion

Despite its importance in plant development, our understanding of the precise structures and physical properties of callose and its function within cell wall matrices is limited. Our study reveals a heterogeneity in plant callose, challenging the prevailing view of callose as a uniform amorphous polymer. We identified two distinct epitopes, using the newly generated LM-BDG1 and LM-BDG2 mAbs and a range of different callose antigens. LM-BDG1 binds specifically to the β-(1,3)-glucan backbone and is therefore an excellent generalised probe for callose detection. LM-BDG2 only binds a subset of β-(1,3)-glucans, different in size and structural conformation. These tools, used in conjunction with chromatographic methodologies and super-resolution microscopy, enabled us to identify, for the first time, callose subpopulations in plants that can be distinguished by size and charge. According to our data, callose interacts with both xyloglucans and pectins forming structurally complex networks of yet unknown function.

The release of callose after digestion of cellulose microfibrils supports previous reports on β-(1,3)-glucan polymer interactions influencing the physicomechanical properties of cellulose hydrogels (*35, 38*). The integration of callose in the cellulose microfibril network was also observed via microscopic observations in forming papillae after fungal infection (*39*) and in poplar expressing *cals3m* (*33*). Our findings indicate the existence of a pool of callose interacting with cellulose microfibrils but also the formation of more structurally complex composites with xyloglucan. The physical binding of xyloglucans to cellulose microfibrils is proposed to regulate cell wall extensibility by creating ‘biomechanical hotspots’(*1*). Our evidence of callose-xyloglucan interactions adds complexity to this model providing another regulatory factor of cell wall properties that requires further investigation. Recent research identified the presence at plasmodesmata of distinct xyloglucans (mostly fucosylated) (*40*) and xyloglucan endotransglucosylases/hydrolases (XTH) involved in xyloglucan metabolism (*41*) providing a biological context, and potential functional significance, for callose-xyloglucan interactions.

Collectively, our findings highlight the structural heterogeneity of plant calloses (we intentionally use the term calloses to capture this heterogeneity) contributing to the properties of distinct microdomains within cell walls. As far as we know, this is the first evidence for the existence of multiple calloses reflecting differences in both the callose polysaccharide *per se* and its interactions with other polysaccharides, both factors underscoring the complexity of cell walls. The use of discriminatory mAbs that can distinguish these structural variations will enable detailed investigations into callose functions in the context of cell wall polymer networks, aiming to disentangle their roles at diverse locations. This in turn will underpin biotechnological approaches to target callose and/or cell walls for a range of applications.

## Acknowledgements

Authors thank Rishikesh Bhalerao for providing the Aspen material. We acknowledge imaging support from the Advanced Bioimaging Laboratory (RRID:SCR_018951) at the Danforth Plant Science Center and usage of the ZEISS Elyra 7 Super-Resolution Microscope acquired through an NSF Major Research Instrumentation grant (DBI-2018962). We are grateful to Fidabio for the loan of a Fida 1 machine.

## Funding

United Kingdom Research and Innovation Future Leader Fellowship MR/T04263X/1 (YBA, RY)

The Leverhulme Trust Grant RPG-2016-136 (YBA, SA)

Biotechnology and Biological Sciences Research Council Discover Fellowship: BB/T009691/1 (SA)

University of Leeds Gosden Studentship (LG)

Engineering and Physical Sciences Research Council Synthetic Biology Postdoctoral Fellowship EP/W022842/1 (JFR)

National Science Foundation MCB 2210127 (TBS)

National Science Foundation NSF-MCB-EAGER 2130365 (KC) ANR-21-CE13-0016-01 DIVCON (EMB)

Human Frontier Research program project RGP0002/2020 (EMB, TSM)

French government in the framework of the IdEX Bordeaux University “Investments for the Future” program / GPR Bordeaux Plant Sciences (EMB).

## Authors contributions

Conceptualization: SA, YBA, PK

Data curation: SA, YBA, JFR, WW, IL, EMB

Formal Analysis: SA, JFR, JB, SEM, WW, IL

Funding acquisition: SA, YBA, PK, EMB, WW, KC, JFR

Investigation: SA, JFR, JB, SEM, IWM, WW, EMB, TSM, YBA

Methodology: SA, YBA, JFR, JB, SEM, IWM, KC, JW, AK, IL, WW, RY, LG, TSM, TBS

Project administration: SA, YBA

Visualization: SA, YBA, JFR, AK, KC, RY, LG, EMB, TSM

Writing – original draft: SA, YBA, JFR, JB

Writing – review & editing: SA, YBA, PK, JFR, KC, WW, EMB, TBS

Supervision: YBA

Software: JFR

## Data and materials availability

All data needed to evaluate the conclusions in the paper are present in the paper and/or the Supplementary Materials. Material will be made available upon request. Raw data are archived at the Research Data Leeds Repository at https://archive.researchdata.leeds.ac.uk/.

## Supplementary Materials

Materials and Methods

Figs. S1 to S8

References (*42-45*)

## References

1. D. J. Cosgrove, Structure and growth of plant cell walls. Nat Rev Mol Cell Biol 25, 340–358 (2024).

2. R. Sinclair et al., Plant cytokinesis and the construction of new cell wall. FEBS Lett 596, 2243–2255 (2022).

3. H. Hofte, A. Voxeur, Plant cell walls. Curr Biol 27, R865–R870 (2017).

4. L. German, R. Yeshvekar, Y. Benitez-Alfonso, Callose metabolism and the regulation of cell walls and plasmodesmata during plant mutualistic and pathogenic interactions. Plant Cell Environ 46, 391–404 (2023).

5. S. Carroll et al., Altering arabinans increases Arabidopsis guard cell flexibility and stomatal opening. Curr Biol 32, 3170–3179 e3174 (2022).

6. K. Jonsson, O. Hamant, R. P. Bhalerao, Plant cell walls as mechanical signaling hubs for morphogenesis. Curr Biol 32, R334–R340 (2022).

7. Y. Gao et al., Elongated galactan side chains mediate cellulose-pectin interactions in engineered Arabidopsis secondary cell walls. Plant J 115, 529–545 (2023).

8. Y. S. Y. Hsieh, M. R. Kao, M. R. Tucker, The knowns and unknowns of callose biosynthesis in terrestrial plants. Carbohydr Res 538, 109103 (2024).

9. P. Kirk, Y. Benitez-Alfonso, Plasmodesmata Structural Components and Their Role in Signaling and Plant Development. Methods Mol Biol 2457, 3–22 (2022).

10. S. Amsbury, P. Kirk, Y. Benitez-Alfonso, Emerging models on the regulation of intercellular transport by plasmodesmata-associated callose. J Exp Bot 69, 105–115 (2017).

11. C. Caseiro, J. N. R. Dias, C. M. G. de Andrade Fontes, P. Bule, From Cancer Therapy to Winemaking: The Molecular Structure and Applications of beta-Glucans and beta-1, 3-Glucanases. Int J Mol Sci 23, (2022).

12. V. Chaudhari et al., Therapeutic and Industrial Applications of Curdlan With Overview on Its Recent Patents. Front Nutr 8, 646988 (2021).

13. R. Hurtado-Guerrero et al., Molecular mechanisms of yeast cell wall glucan remodeling. J Biol Chem 284, 8461–8469 (2009).

14. K. Kapoor, A. Geitmann, Pollen tube invasive growth is promoted by callose. Plant Reprod 36, 157–171 (2023).

15. J. Liu et al., APP1/NTL9-CalS8 module ensures proper phloem differentiation by stabilizing callose accumulation and symplastic communication. New Phytol 242, 154–169 (2024).

16. E. M. Bayer, Y. Benitez-Alfonso, Plasmodesmata: Channels Under Pressure. Annu Rev Plant Biol, (2024).

17. A. Vaten et al., Callose biosynthesis regulates symplastic trafficking during root development. Dev Cell 21, 1144–1155 (2011).

18. S. W. Wu, R. Kumar, A. B. B. Iswanto, J. Y. Kim, Callose balancing at plasmodesmata. J Exp Bot 69, 5325–5339 (2018).

19. A. F. Sankoh, T. M. Burch-Smith, Plasmodesmata and hormones: pathways for plant development. Am J Bot 108, 1580–1583 (2021).

20. E. E. Tee, C. Faulkner, Plasmodesmata and intercellular molecular traffic control. New Phytol, (2024).

21. S. Amsbury, Y. Benitez-Alfonso, Immunofluorescence Detection of Callose in Plant Tissue Sections. Methods Mol Biol 2457, 167–176 (2022).

22. A. F. Sankoh, J. Adjei, D. M. Roberts, T. M. Burch-Smith, Comparing Methods for Detection and Quantification of Plasmodesmal Callose in Nicotiana benthamiana Leaves During Defense Responses. Mol Plant Microbe Interact, MPMI09230152SC (2024).

23. V. Cornuault, I. W. Manfield, M. C. Ralet, J. P. Knox, Epitope detection chromatography: a method to dissect the structural heterogeneity and inter-connections of plant cell-wall matrix glycans. Plant J 78, 715–722 (2014).

24. Y. Verhertbruggen, S. E. Marcus, A. Haeger, J. J. Ordaz-Ortiz, J. P. Knox, An extended set of monoclonal antibodies to pectic homogalacturonan. Carbohydr Res 344, 1858–1862 (2009).

25. S. Vidal-Melgosa et al., A new versatile microarray-based method for high throughput screening of carbohydrate-active enzymes. J Biol Chem 290, 9020–9036 (2015).

26. H. R. Johnsen et al., Cell wall composition profiling of parasitic giant dodder (Cuscuta reflexa) and its hosts: a priori differences and induced changes. New Phytol 207, 805–816 (2015).

27. M. Mirdita et al., ColabFold: making protein folding accessible to all. Nat Methods 19, 679–682 (2022).

28. J. Jumper et al., Highly accurate protein structure prediction with AlphaFold. Nature 596, 583–589 (2021).

29. A. K. Nivedha, D. F. Thieker, S. Makeneni, H. Hu, R. J. Woods, Vina-Carb: Improving Glycosidic Angles during Carbohydrate Docking. J Chem Theory Comput 12, 892–901 (2016).

30. S. Becker et al., Laminarin is a major molecule in the marine carbon cycle. Proc Natl Acad Sci U S A 117, 6599–6607 (2020).

31. H. Saito, A. Misaki, T. Harada, A Comparison of the Structure of Curdlan and Pachyman. Agricultural and Biological Chemistry 32, 1261–1269 (1968).

32. B. K. Patel, O. H. Campanella, S. Janaswamy, Impact of urea on the three-dimensional structure, viscoelastic and thermal behavior of iota-carrageenan. Carbohydr Polym 92, 1873–1879 (2013).

33. M. Bourdon et al., Ectopic callose deposition into woody biomass modulates the nano-architecture of macrofibrils. Nat Plants 9, 1530–1546 (2023).

34. C. Simpson, C. Thomas, K. Findlay, E. Bayer, A. J. Maule, An Arabidopsis GPI-anchor plasmodesmal neck protein with callose binding activity and potential to regulate cell-to-cell trafficking. Plant Cell 21, 581–594 (2009).

35. R. H. Abou-Saleh et al., Interactions between callose and cellulose revealed through the analysis of biopolymer mixtures. Nat Commun 9, 4538 (2018).

36. H. L. Pedersen et al., Versatile high resolution oligosaccharide microarrays for plant glycobiology and cell wall research. J Biol Chem 287, 39429–39438 (2012).

37. V. Cornuault, J. P. Knox, Sandwich Enzyme-linked Immunosorbent Assay (ELISA) Analysis of Plant Cell Wall Glycan Connections. Bio-protocol 4, e1106 (2014).

38. P. Kumari, P. Ballone, C. Paniagua, R. H. Abou-Saleh, Y. Benitez-Alfonso, Cellulose-Callose Hydrogels: Computational Exploration of Their Nanostructure and Mechanical Properties. Biomacromolecules 25, 1989–2006 (2024).

39. D. Eggert, M. Naumann, R. Reimer, C. A. Voigt, Nanoscale glucan polymer network causes pathogen resistance. Sci Rep 4, 4159 (2014).

40. A. Paterlini et al., Enzymatic fingerprinting reveals specific xyloglucan and pectin signatures in the cell wall purified with primary plasmodesmata. Front Plant Sci 13, 1020506 (2022).

41. S. Gombos et al., A high-confidence Physcomitrium patens plasmodesmata proteome by iterative scoring and validation reveals diversification of cell wall proteins during evolution. New Phytol 238, 637–653 (2023).

42. A. Lees, B. L. Nelson, J. J. Mond, Activation of soluble polysaccharides with 1-cyano-4-dimethylaminopyridinium tetrafluoroborate for use in protein-polysaccharide conjugate vaccines and immunological reagents. Vaccine 14, 190–198 (1996).

43. I. Moller et al., High-throughput screening of monoclonal antibodies against plant cell wall glycans by hierarchical clustering of their carbohydrate microarray binding profiles. Glycoconj J 25, 37–48 (2008).

44. G. M. Morris et al., AutoDock4 and AutoDockTools4: Automated docking with selective receptor flexibility. J Comput Chem 30, 2785–2791 (2009).

45. J. Walton, Lead aspartate, an en bloc contrast stain particularly useful for ultrastructural enzymology. J Histochem Cytochem 27, 1337–1342 (1979).

