## Supplemental Methods and figures for "Structurally diverse calloses/β-1,3-glucans in plant cell wall microdomains"

**The PDF file includes:**

Materials and Methods
Figs. S1 to S8
References

### Materials and Methods

Plant materials

*Arabidopsis thaliana* ecotype Columbia Col-0, and transgenic lines *cals3m* (a pG1090- estradiol inducible version of the mutant CAL3 described in (*17, 35*)) and PDCB1-OE (pCaMV*35S*-PDCB1-YFP described in (*34*)) were used. Seeds were surface-sterilized in bleach solution (20% thin bleach, 0.01% tween-20) for 10 min and stratified in the dark at 4 °C for 5 days. Seeds were germinated on ½ Murashige and Skoog (MS) medium plates containing 0.22% MS basal medium, 1% sucrose, 0.8% plant agar and grown under long day conditions (16 h photoperiod) at a constant temperature of 20 °C. For *cals3m* induction, ten day old pG1090::i*cals3m* seedlings were transferred for induction to ½ MS plates supplemented with 10 µM β-estradiol (Sigma Aldrich, diluted from a stock solution in DMSO). Control experiments (with basal/non-induced amount of callose) were performed by exposing pG1090::i*cals3m* to the corresponding solvent (DMSO) concentration or by transferring wild type seedlings to 10 µM β-estradiol.
For tomato work *Solanum lycopersicum* variety Ailsa Craig plants were grown in compost (John Innes No. 1, UK) in 30 $cm^2$ round pots. The plants were grown in a glasshouse at 25°C, 16h photoperiod, at 60% relative humidity. Fruits were harvested at 4 weeks post anthesis. The pericarp was cut into ~1$cm^2$ pieces that were fixed in 4% Paraformaldehyde (Generon, UK) as described below.
*Nicotiana benthamiana* seeds were sterilised using 70 % ethanol for 10 mins, with constant shaking. Seeds were washed in 100 % ethanol before stratifying in ½ MS plates for 4 days at 4 °C. Plants were germinated in long day light conditions (16-hour day) at 22°C and grown for 8 weeks in sand watered twice a week with 1x Long Ashton nutrient media. Roots were collected, washed and fixed in paraformaldehyde as described for tomato above.
Hybrid aspen buds (*Populus tremula x tremuloides*) clone T89 (wild type) were collected after 11 weeks of growth in short photoperiod (8-hour, 20°C light/16-hour, 15°C dark cycles) and fixed as above.

Polysaccharide materials

Curdlan (Megazyme), Pachyman (Megazyme), laminarin (Sigma) and other glycan/ polysaccharide standards were acquired from the commercial suppliers and dissolved following the supplier's protocols. oligo-β-(1,3)-gluco-saccharides from a disaccharide (laminaribiose) to a hexosaccharide (laminarihexaose) were obtained from Megazyme. For screening of purified polysaccharides, 1 mg/ml (w/v) of samples were dissolved either in 0.01 M PBS, pH 7.4 or 1 M KOH (for those insoluble in PBS such as pachyman, curdlan and scleroglucan).
FITC-dextran used as standards for size exclusion chromatography were obtained from Sigma and prepared as suggested by the suppliers. A Fida 1 system (https://www.fidabio.com/) was used for hydrodynamic analysis of FITC-dextrans using fluorescence detection with excitation at 480 nm.

Isolation of monoclonal antibodies.

Antibodies were produced as previously described (*24*). Laminarin was coupled to ovalbumin using the CDAP method (*42*). The conjugate in Complete Freunds Adjuvant was administered subcutaneously at Day 0 and Freunds incomplete adjuvant on Day 25. The immune response was gauged by testing binding of blood serum to pachyman 10 days after the second injection. A prefusion intraperitoneal injection of the conjugate without adjuvant was administered on Day

66. Spleen lymphocytes were then isolated 3 days later and fused with rat myeloma cell line IR983F. Viable antibody cell lines were selected by ELISA using 50 ug/ml Pachyman as the immobilised antigen. The cell lines of interest were cloned to produce the monoclonal antibodies LM-BDG1, LM-BDG2 and LM-BDG3. Immunoglobulin isotyping was carried out with LM-BDG1 and LM-BDG2 being IgM and LM-BDG2 being IgG2c. Sequencing of LM-BDG1 and LM-BDG2 variable regions was carried out by Absolute Antibody (https://absoluteantibody.com/).
LM19 and LM25 antibodies were produced and obtained by Plant Probes (https://plantcellwalls.leeds.ac.uk/science/antibodies/). (1-3)-beta-glucan-directed monoclonal antibody BioS (400-2) was obtained from Biosupplies (http://www.biosupplies.com.au/) and prepared following provider's protocols.

Cell wall extractions

To prepare alcohol insoluble residues (AIR), leaf material was flash frozen and freeze-dried before being weighted and transferred to an eppendorf with two 3 mm stainless steel ball bearings. The material was ground to a fine powder in a Qiagen TissueLyserII (Qiagen, Hilden, Germany) at 30 Hz for 1 min. 250 mg of ground tissue was sequentially incubated in 1 ml volumes of a solvent series consisting of ethanol (50%, 60%, 70%, 80%, 90% and 100% v/v) followed by acetone and a chloroform:methanol mixture (3:1). Each stage samples were incubated for 90 minutes on a rocking table at room temperature, samples were pelleted by centrifugation and the solvent discarded. Following the final step, the sample was dried by evaporation leaving AIR which is enriched in cell walls.
2 mg of AIR was incubated with 1 ml of 50 mM cyclohexanediaminetetraacetic acid (CDTA), pH 6 for 90 minutes and shook at 10 Hz in a Qiagen TissueLyserII. Undissolved sample was pelleted by centrifugation and the supernatant retained as the CDTA extract. This extraction process was repeated with 4 M potassium hydroxide and retained as the KOH extract. Any remaining residues was washed and subjected to digestion by incubating for 8 h at 30°C with 1 mg/ml of cellulase 5A (NZYTech, Lisbon, Portugal) in 20 mM Tris-HCl buffer pH 8.8 to give the cellulase extract. The KOH extract was neutralised to pH 7, using 80% (v/v) acetic acid, immediately prior to use.

Glycomicroarrays

A set of well-defined glycans, including ß-1,3 and ß-(1,3-1,4)-glucans, were dissolved in 18.2 MΩ/cm $H_2O$ to a final concentration of 1 mg/mL and placed on a rotary shaker (Cole-Parmer, St Neots, UK) at 4 °C for 12 hours to ensure complete solubilisation. All samples were added to wells of a 384-well microtiter plate (Greiner Bio-One, Kremsmünster Austria) and diluted 1:1 (v/v) with printing buffer (47 % glycerol, 52.9 % deionised water, 0.06 % Triton X-100 and 0.04 % ProClin™ 200) to a final volume of 40 µL in each well. Glycomicroarrays were prepared as previously described by Vidal-Melgosa et al. (*25*) and were printed onto nitrocellulose membrane with a Marathon Argus Microarray Printer (Arrayjet, Roslin, UK). Probing of printed microarrays was performed according to Johnsen et al. (2015) with slight modifications. Briefly, microarrays were blocked for 2 hours at room temperature in PBS supplemented with 5 % (w/v) skimmed milk powder (MP/PBS). Following incubation, arrays were probed for 2 hours with primary antibodies. Arrays were rinsed thoroughly with clean PBS and then incubated for 2 h with secondary antibody anti-rat or anti-mouse (as appropriate) IgG coupled to alkaline phosphatase, diluted 1:1000 in MP/PBS. Arrays were rinsed extensively with clean PBS following incubation and binding of mAbs to the nitrocellulose membrane was detected by 5-bromo-4-chloro-3'-indolyphosphate p-toluidine (BCIP) salt and nitro-blue tetrazolium (NBT) chloride. Microarrays were then submerged in fresh $H_2O$ and allowed to dry overnight between filter papers. Microarray probing was performed in triplicate for each probe and control. Developed microarrays were scanned at 2400 dpi (Canon CanoScan 8800F, Denmark) and the images saved as negative, 16-bit TIFF files. Binding intensities of probes to individual spots was quantified using microarray analysis software (Array-Pro Analyzer version 6.3) fitted with an automated grid tool. A value of 100 was assigned to the highest mean spot signal intensity recorded and all other values were normalised accordingly to allow quantification of the relative glycan epitope abundance for each sample (*43*).

Size exclusion chromatography (SEC).

Cell wall extracts were diluted 1:5 in 20 mM sodium acetate, 1 M sodium chloride (running buffer) and separated using a HiPrep 16/60 sephacryl S-400 HR size exclusion column (Cytiva life sciences, UK) and an ÄKTA explorer system. Samples were eluted at 1 ml per minute with running buffer, 25ml was discarded as void volume followed by collection of 96 x 1 ml fractions. 40 µl of 1M sodium carbonate was added to each eluted fraction. For SEC of individual polysaccharides, samples were diluted to 50 µg/ml in running buffer and ran as described above.

Anion exchange chromatography (AEC)

Chromatography was carried out as described by (*23*). Cell wall extracts were diluted 1:5 in 20 mM sodium acetate (low salt buffer) and separated using a HiTrap ANX FF (High Sub) anion exchange column (Cytiva life sciences, UK) using an ÄKTA explorer system. Samples were eluted at 1 ml per minute with Low salt buffer for 35ml, followed by mixing of low salt buffer with high salt buffer (50 mM sodium acetate, 1 M NaCl) to create an increasing salt gradient ranging from 0% high salt buffer (at 35 ml) to 100% high salt buffer (at 60 ml) followed by further elution in 100% high salt buffer until a total of 96 x1 ml fractions were collected. 40 µl of 1M sodium carbonate was added to each eluted fraction.

ELISA

Polysaccharide samples were diluted to 50 µg/ml in PBS and used to coat an immunosorp 96-well plates (Maxisorp, F96, Thermofisher), 100 µl per well, and incubated overnight at 4 °C. SEC fractions of plant extracts or polysaccharides were coated directly on to plates using 100 µl per well of the fraction. After coating, plates were rinsed in tap water and blocked for 1 h at room temperature using 200 µL PBS containing 5% (w/v) non-fat bovine milk powder (Sigma) followed by extensive washing with tap water. 100 µL per well of primary antibody solution (LM-BDG1/2 hybridoma supernatant diluted 10x or BioS diluted 100x in PBS containing 5% (w/v) milk powder) was added and incubated for 2 h at room temperature. After incubation, plates were washed with tap water, and 100 µL per well of anti-rat IgG –HRP (Thermofisher Catalog # 31470) or anti-mouse IgG-HRP (Catalog # 31430) for BioS, at 1000-fold dilution in PBS containing 5% (w/v) milk powder, was added and incubated for 1 h followed by another washing step. The plates were developed by adding 100 µL per well of substrate solution (to make 20 ml of substrate 2 ml of 1M sodium acetate pH 6; 200µl of 10 mg/ml 3,3',5,5'-tetramethylbenzidine in DMSO and 20 µl of 6% hydrogen peroxide were added to 17.78 ml of water) and incubating for 6 min. The reaction was stopped by adding 50 µL per well of 2.5 M

$H_2SO_4$, resulting in the formation of a yellow colour which was measured as absorbance at 450 nm in a plate reader. ELISA data was baselined by subtracting negative controls (PBS coated wells) from the absorbance readings, negative values were corrected to 0. Graphs were constructed using Graphpad PRISM software.

Hapten inhibition assay

To determine working antibody concentration, ELISA was carried out as described above using curdlan coated plates and a dilution series of primary antibody from 1x dilution to 2500 x dilution. For the hapten inhibition assay, a competitive inhibition ELISA was conducted. LM-BDG1 and LM-BDG2 were used at an antibody dilution that gave 90% of maximal binding. Plates were coated overnight with 100 µl of 50 µg/ml curdlan. After washing, 50 µl per well of serially diluted hapten (oligosaccharides diluted from 1000 µg/ml (w/v) to 0.064 µg/ml (w/v) in PBS) was added. 50 µl per well of primary antibody (LM-BDG1 and LM-BDG2) were added to the haptens at 2x the working concentration determined above and incubated for 90 minutes with agitation. Plates were washed extensively in tap water and the rest of the ELISA carried out as described above. Primary antibodies bound to haptens will be removed (during the washing) and only when bound to curdlan will be retained which produces the ELISA signal.

Urea assay

1 mg/ml (w/v) curdlan samples were diluted to a final concentration of 50 µg/ml (w/v) in urea dissolved in PBS. Urea was used at the following concentrations: 4, 3, 2, 1, 0.5, 0.1, 0 M. Curdlan:Urea mixtures were then coated onto plates overnight and ELISA carried out as described above.

LM-BDG1 purification and conjugation with HRP

50 mL hybridoma supernatant was filtered using a 0.45 µm filter. The filtered hybridoma supernatant was diluted 1:1 with Binding Buffer (100 mM sodium phosphate, 150 mM sodium chloride, pH 7.2) and applied at a rate of 1 ml/min to a 1 mL protein L chromatography cartridge (Pierce, ThermoFisher) equilibrated with 10 mL of Binding Buffer. The cartridge was washed with 10 mL Binding Buffer, until the $A_{280}$ returned to baseline level, indicating removal of unbound proteins. LM-BDG1 was then eluted with 5 mL Elution Buffer (0.1 M glycine, pH 2.8). Elution fractions of 1 mL were collected in tubes containing 0.1 mL Neutralization Buffer (1 M Tris-HCl, pH 8.5). Fractions containing protein (indicated by $A_{280}$) were pooled and the pooled fractions buffer exchanged into PBS and concentrated using a centrifugal filter (Amicon Ultra-15, 50 kDa MWCO, Merck) and checked for purity using reduced SDS-PAGE. 5 µL LM-BDG1 was mixed with 1.5 µL ultrapure water, 2.5 µL 4 × LDS sample buffer (NuPAGE, ThermoFisher) and 1 µL 10 × reducing agent (NuPAGE, ThermoFisher), and heated to 95 °C for 5 min. The sample was run on a 12 % Tris-Glycine precast acrylamide gel (NuSep) at 200 V for 30 min, in 1 × Tris-Glycine-SDS Running Buffer (25 mM tris base, 200 mM glycine, 0.1 % *w/v* SDS). The gel was briefly rinsed in ultrapure water and stained in Brilliant Blue R stain solution (3 mM Brilliant Blue R, 45 % *v/v* methanol, 10 % *v/v* acetic acid) for 1 h. The gel was destained in destaining solution (45 % *v/v* methanol, 10 % *v/v* acetic acid) for 2 h and rinsed with water, before imaging. Purified LM-BDG1 was conjugated to horseradish peroxidase (HRP) using the LYNX rapid HRP conjugation kit (LNK006P, BioRad), as specified by the manufacturer.

Sandwich ELISA

Sandwich ELISA was carried out as described by (*37*). Monoclonal antibodies (hybridoma supernatants) or purified CBM43 (*34*) were diluted 10 times in PBS and 100 µL per well was used to coat 96-well immunoplates (Maxisorp, F96, ThermoFisher). PBS was added to no-coated control wells. After overnight incubation at 4 °C, wells were washed 3 times with 300 µL per well of 0.05 % (v/v) Tween-20 in PBS. To block uncoated spaces, plates were incubated with gentle rocking for 2 h at room temperature with 300 µL per well 5 % (w/v) milk powder in PBS. Cell wall extracts were diluted 10 times and 100 times in 5 % (w/v) milk in PBS. Pachyman 1,3-β-D-glucan (Megazyme) was diluted to 100 pg/mL in 5 % (w/v) milk in PBS, for the control. Plates were washed as before and incubated with 100 µL per well diluted cell wall extract or Pachyman, for 1 h with gentle rocking at room temperature. Purified LM-BDG1 labelled with HRP was diluted 5000 times in 5 % (w/v) milk in PBS. Plates were washed and incubated for 1 h with gentle rocking, at room temperature, with 100 µL per well diluted LM-BDG1-HRP. Plates were washed and incubated with 100 µL per well 3,3',5,5'-tetramethylbenzidine (TMB chromogen solution, ThermoFisher) for 10 min at room temperature. The reaction was stopped by adding 50 µL per well 2.5 M sulfuric acid. Absorbance was read at 450 nm.

Molecular docking analysis.

Antibody models were predicted from sequencing data using the Sokrypton ColabFold version of AlphaFold (AlphaFold2_mmseqs2 (*27, 28*), including heavy and light chains, and both constant and variable regions. Due to the Fc region being absent from the prediction, there was a reduction in pLDDT at the Fab to Fc interface and a minor reduction in pLDDT at the ligand binding cleft, (dipping below pLDDT of 90 for stretches of 5 amino acids and reaching a minimum pLDDT of 78.2 for one amino acid) still within the 'confident' prediction range. The variable region of the top scoring pLDDT models were then used for docking with Vina-Carb. Five models of beta-1-3-glucan polysaccharides were generated in PyMOL using the beta 1,3 glucan pentasaccharide retrieved from PDB:2w62 (*13*). This pentasaccharide was used to construct the 'glucan-2' to 'glucan-6' polysaccharides by truncating or extending the pentasaccharide respectively in PyMOL. Molecular docking was then performed on LM-BDG1 and LM-BDG2 with each of the 5 different length glucans. For simplicity the constant region was removed before docking. In each case residues were manually identified in the binding cleft for 'flexible docking' which allows rotameric side chain conformations to be sampled during the docking. PDBQT files and the grid file were prepared on AutoDock Tools (ADT, (*44*)). Vina-Carb (*29*) was used to generate ten ensembles of nine docked poses each for the range of 2 to 6 residue beta-1-3-glucans. All reported energies are chi adjusted affinities, accounting for glycosidic bond linkages. RMSD based energy landscapes were generated via an in-house script. RMSD values between glucans were determined with PyMOL, and then metricised, clustered and visualised with python packages sklearn, scipy and seaborn, respectively, among others.

RMSD energy landscapes for docked glucans

For each antibody (LM-BDG1 and LM-BDG2), 1-3 beta-glucan oligomers of 2, 3, 4, 5 and 6 residues were docked with Vina-Carb. A 2D RMSD matrix between each pair of bound beta-glucans was generated in PyMOL. This matrix was symmetrically clustered via agglomerative clustering in scikit-learn (python) and grouped via hierarchy linkage in scipy (python). The interaction energy calculated by vina-carb was averaged over each hierarchy grouping and superimposed on top of the 2D RMSD clustered matrix to give an RMSD based interaction energy landscape. From this landscape, low-energy, high-population wells were identified. The lowest interaction energy member of this group was identified as the prospective most likely binding position and orientation.

Preparation of plant samples for microscopy

Plant material was harvested and immediately fixed in 4% (w/v) formaldehyde in PEM buffer (0.1 M PIPES, 2 mM EGTA, 1 mM MgSO4, adjusted to pH 7) by vacuum infiltration. After fixation, both Arabidopsis and Tomato samples were dehydrated in an ethanol series (30 min each at 30%, 50%, 70%, 100% EtOH) and infiltrated in Steedmans wax (polyethylene glycol 400 distearate and 1-hexadecanol in a 9:1 ratio) by sequential incubation in paraffin wax and ethanol, (1:3, 1:1, 3:1 v/v) for 4 hours each followed by 100% wax (2 x 1 h). Samples were then embedded in 100% Steedmans wax which was allowed to harden for 5 h at room temperature. Wax sections were cut to a thickness of 9 µm using a HM 325 rotary microtome (Microm, Bicester, UK) and transferred to polylysine coated microscope slides. Wax was removed by incubation in 100% ethanol for 3x 10 minutes and rehydrated by a descending ethanol:water series: 90% (v/v), 50% (v/v), water (10 min per change) followed by a final incubation for 90 min in water. Slides were then air dried and used for microscopy.

In the case of tobacco small, fixed root segments were sandwiched gently in the middle of Elder wood cylinders (supplied from Eternal Tools) cut half longitudinally. Wilkinson sword, Double Edge Blades were used to finely cut the root transversally. Once resuspended in deionised water, the sections were selected based on their visible thickness under light microscope and moved to 96-well plates for immunolocalization.

All other samples used for immunofluorescence (except $SIM^2$ labelling as described below) were infiltrated with LR White Resin (London Resin Company) diluted in ethanol (45 min each at 10%, 20%, 30%, 50%, 70% & 90% resin then 3x8 h at 100%). Samples were then inserted in gelatine capsules filled with resin and allowed to polymerize for 7 days at 37°C. Sections were cut to a thickness of 2 µm using a Reichert-Jung Ultracut E ultramicrotome and a glass knife.

Sample preparation for immunogold and immunofluorescence $SIM^2$ labelling

Tobacco leaf pieces were fixed in a freshly prepared solution of 2% paraformaldehyde (Cat. No. 15710, Electron Microscopy Sciences, Hatfield, PA, USA), 2% glutaraldehyde (Cat. No. 16020, EMS, Hatfield, PA, USA), 0.01% Tween20 (Cat. No. 20605, USB Corporation, Cleveland, OH, USA), 0.05% malachite green (Cat. No. M9015-25G, Sigma-Aldrich, St. Louis, MO, USA) in 0.1M sodium cacodylate buffer (pH 7.4) (Cat. No. 11653, EMS, Hatfield, PA, USA) by vacuum infiltration for 30 minutes followed by overnight incubation at 4 °C. Samples were postfixed in 2% osmium tetroxide (Cat. No. 19110, EMS, Hatfield, PA, USA) for 4 h at room temperature and rinsed 3 times with distilled water. Subsequently, samples were *en bloc* stained with 1% aqueous uranyl acetate (Cat. No. 02624-AB, SPI Supplies, West Chester, PA, USA) (overnight in 4 °C followed by incubation in the oven at 50 °C for 2 h) and freshly prepared lead aspartate (50 °C for 2 h) (*45*) and rinsed 3 times with distilled water after each staining step. Samples were then dehydrated using a graded cold acetone series of 25%, 50%, 75%, 95%, 2x 100%, 30 mins each) (Cat. No. 10015, EMS, Hatfield, PA, USA) and treated with 100% propylene oxide (Cat. No. 20401, EMS, Hatfield, PA, USA) (2 times, 30 mins) to ensure consistent resin infiltration. Samples were infiltrated with propylene oxide:Quetol 651/NSA resin (Cat. No. 14640, EMS, Hatfield, PA, USA) (without the DMP-30 catalyst) at 3:1, 2:1, 1:1, 1:2, 1:3 and 2x 100% Quetol for 6-12 h each. Subsequently, two changes of 100% Quetol 651/NSA with the DMP-30 catalyst

were performed and samples were embedded in Quetol with DMP30 at 60 °C for 48 h. Embedded samples were then trimmed with a razor blade and sectioned using a Leica Ultracut UCT ultramicrotome with a Diatome diamond knife. Sections for immunogold labelling (70 nm) were collected onto gold formvar-carbon cotted single-slot grids and sections for immunofluorescence labelling (200 nm) were collected on to glass coverslips.
Aspen buds and Arabidopsis cell culture were fixed in 2% glutaraldehyde and 2% paraformaldehyde in 0.02 M cacodylate buffer (pH 7.2), combined with 1% tannic acid; postfixed 1% $OsO_4$ in water, dehydrated, infiltrated and embedded in Spurr's resin (Ted Pella). The blocks were sectioned longitudinally into slices with a thickness of 90 nm using an EM UC7 ultramicrotome (Leica). These sections were then delicately positioned onto 100 mesh copper grids and immunogold labelling was performed (see below). Subsequent observations were conducted employing a FEI TECNAI Spirit 120 kV electron microscope.

Immunofluorescence labelling

For immunofluorescence of callose, plant sections were incubated in 0.1 M Na2CO3, pH 11.4 for 1 hour to remove methyl side groups from pectin. During each incubation samples were gently rocked at 25 degrees, after which the sections were washed 3-4 times with PBS. Following this the pectin was digested by pectate lyase 10 ug/ml in 0.1 M CAPS buffer for 4 hours. Blocking was achieved using 3% (w/v) milk protein in PBS (PBS/MP). Sections were then incubated with a ten-fold dilution of primary mAb in PBS/MP for 1 h at room temperature. Samples were washed 3 times with PBS and secondary antibody was added (anti-rat-IgG coupled to Alexafluor-488 (Thermofisher, Cat # A-11006) was used at 100-fold dilution in PBS/MP) for 1 h. Samples were kept in the dark from this step. Samples were counterstained with 0.25% (w/v) calcofluor white or pontamine red solution diluted ten-fold in PBS for 5 min before washing 3 times with PBS and mounting on slides with Citifluor AF1 anti-fade solution (Agar Scientific, UK). Most images were captured using a DP51 camera. FITC was visualized using a filter set with 460-490 nm excitation filter, a 510-550 nm emission filter and a 505 nm dichroic mirror. Calcofluor White was visualized using a 395 nm excitation filter, a 460 nm emission filter and a 425 nm dichroic mirror. Immunofluorescence images were taken on a Zeiss LSM880 upright confocal microscope excited by an Argon laser at 488 nm. Pontamine red was detected at 560–605 nm.

$SIM^2$ Immunofluorescence labelling

Plant sections were attached to Vectabond®-coated (Vector Laboratories, Newark, CA, USA) coverslips No. 1.5 (Cat. No. 72204-10, Electron Microscopy Sciences, Hatfield, PA, USA) following the manufacturer's protocol. Prior to labelling, sections underwent an etching step. An etching solution of 50% sodium methoxide was prepared by diluting saturated sodium methoxide with methanol. Sections on the coverslips were deplasticized by adding the etching solution for 10 minutes followed by rinsing in 1:1 xylene:methanol for 3 minutes and in 100% methanol for 3 minutes. Sections were rehydrated in 1X PBS for 15 minutes. Antigen retrieval was performed by placing the coverslips with sections in 10mM sodium citrate buffer for 10 minutes at 95 °C. Sections were allowed to cool down to RT. For the blocking step, sections were treated with a blocking buffer consisting of 2% bovine serum albumin (BSA) (Cat. No. A9205, Sigma Aldrich) in Pierce protein-free buffer (Cat. No. 37572, ThermoFisher Scientific) for 1 h at room temperature (RT). Three primary antibodies were used for immunofluorescence and immunogold labelling: Rat LM-BDG2, Rat LM-BDG2, Rat non-immune IgG 0.1 mg/mL as a negative control

and showed no specific pattern at cell walls or plasmodesmata. Coverslips were incubated with primary antibodies (20X dilution in blocking buffer) for 2 h at RT inside a humid chamber and rinsed with blocking buffer 3 times. Subsequently, coverslips were incubated with secondary antibody (diluted 1:250 in a blocking buffer, chicken anti-rat DY Light 550 Cat No. AS12 1973, Agrisera, Vannas, Sweden) and Calcofluor White (5mM stock diluted 1:100 Cat. No. 29067, Biotium, Fremont, CA, USA) for 1 h at RT. Coverslips were rinsed with 1x PBS three times and mounted on glass slides and edges were temporarily sealed using Twinsil® (Cat. No. 32002.42011-12, Picodent, Reagent A, Reagent B Cat. No. 34001.02014-06 Wipperfurth, Germany), mixing two components in 1:1 ratio. After the first antibody super-resolution imaging round (e.g., LM-BDG1), the labelled sections were stripped, and coverslips with the same sections were relabelled and for a second round with the second callose antibody (e.g., LM-BDG2) (also labelled in reverse order in separate experiments). For antibody stripping, coverslips with sections-side facing up were incubated in Glycine-SDS buffer (20mM glycine, 10% v/v SDS, adjust pH with HCl to pH 2.0) for 3 hours at 50°C on a shaker.
For super-resolution structure illumination microscopy (SIM), we acquired single-plane or Z stack images, arranged in a tile-scan, using Zeiss Elyra 7 microscope, equipped sCMOS pco.edge camera, and C-Apochromat 63x/1.2 W Korr M27 or C-Apochromat 40x/1.20 W Corr FCS objectives. Raw images were acquired as 1024X1024 (63X) or 1280X1280 (40X) and 16-bit monochrome using Lattice SIM mode with 13 phases and 36.5 µm (63X) or 23.0 µm (40X) grating grid period with 60nm (63X) or 99nm (40X) pixel size. Calcofluor white was excited with 405-nm and DY Light 550 was excited with 561-nm lasers. Emission from each dye was collected sequentially, using frame fast switching mode and the following filters: LBF 405/488//561/642 reflector revolver; BF570-620 + LP655 and BP420-480 + BP495-550 emission filters; SBS LP560 beam splitter. All raw images were then processed in Zeiss Zen 3.0 SR FP2 (black edition) v 16.0 software using the following parameters: $SIM^2$ 2D Leap in Standard-Fixed mode, input SNR set to "Low", iterations set to "20", the regularization weight "0.04" (Calcofluor White) and "Low", iterations set to "22", the regularization weight "0.015" (DyLight 550), processing and output sampling "2" and selected to keep raw data intensity values "Scale to raw." After $SIM^2$ processing, we aligned mismatched channels using a multi-color beads calibration slide (Zeiss, Cat No. 000000-2076-515, Germany) and processing tool "Channel alignment" in Zen Black software. Finally, in some instances, separate images in a tile-scan were also stitched, using "Stitching" tool set to a 0.9 strict correlation threshold. Sequentially acquired multiplex images were aligned using the Calcofluor White stained cell walls to register sections between rounds. Images were opened in Adobe Photoshop CC, overlaid with Linear Dodge (Add), and using the Transform rotate and move tool until the cell walls were registered.

Immunogold labelling

Grids were placed in mPrep/g capsules (Cat. No. 21505, Microscopy Innovations, Marshfield, WI, USA) and the capsules were moved thorough solutions for immunogold labelling. Saturated sodium methoxide was diluted to 10% with methanol for deplasticizing sections. Sections were etched with this solution for 2 minutes, followed by rinsing in 100% methanol twice for 5 minutes, in 10% methanol for 5 minutes and in distilled water 3 times for 5 minutes. Grids were incubated in a blocking buffer of 3% BSA (Cat. No. A9205, Sigma Aldrich, Saint Louis, MO, USA) in Pierce protein free buffer (Cat. No. 37572, ThermoFisher Scientific) and for 1 h at RT. Grids were incubated with a twenty-fold dilution of primary antibodies (described above) in blocking buffer for 2 h at RT and rinsed 5 times with blocking buffer. Then, grids were incubated with a forty-fold dilution of secondary antibody (Cat. No. 112-205-143, 12 nm Colloidal Gold-Affinity Pure Goat Anti-Rat IgG (H+L), Jackson ImmunoResearch Laboratories) in blocking buffer for 1 h at RT and rinsed 5 times with blocking buffer. Subsequently, the grids were incubated in 1% glutaraldehyde for 2 minutes, rinsed 5 times with double-distilled water and allowed to air-dry. Grids were imaged using a ThermoFisher Scientific Talos L120C transmission electron microscope (TEM) at 120kV and high-resolution images were acquired with a Ceta 16M CMOS 4k x 4k camera.

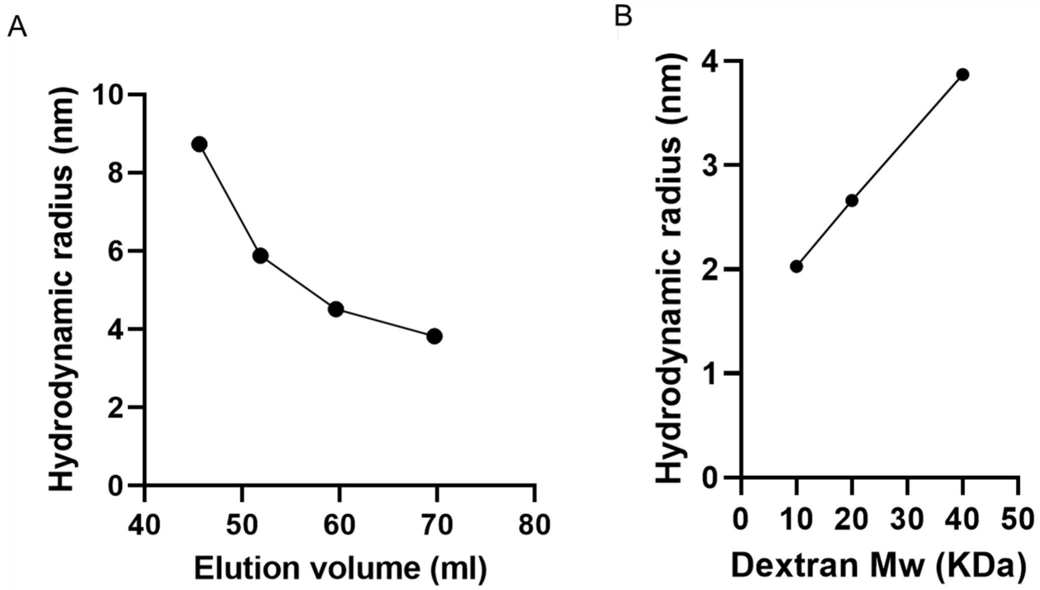


**Fig. S1. Size exclusion separation of standards. (A)** Size exclusion chromatography was carried out using thyroglobulin, ferritin, aldolase and conalbumin protein standards (669, 440, 158 and 75 kDa respectively). Elution volume vs hydrodynamic radius (determined using FIDA) is represented. **(B)** The hydrodynamic radius of dextran standards were calculated using FIDA and plotted against their MW. A 40 kDa dextran has a hydrodynamic radius of 3.87 nm equivalent to a protein with elution volume of ~70 ml and similar MW as Pachyman.

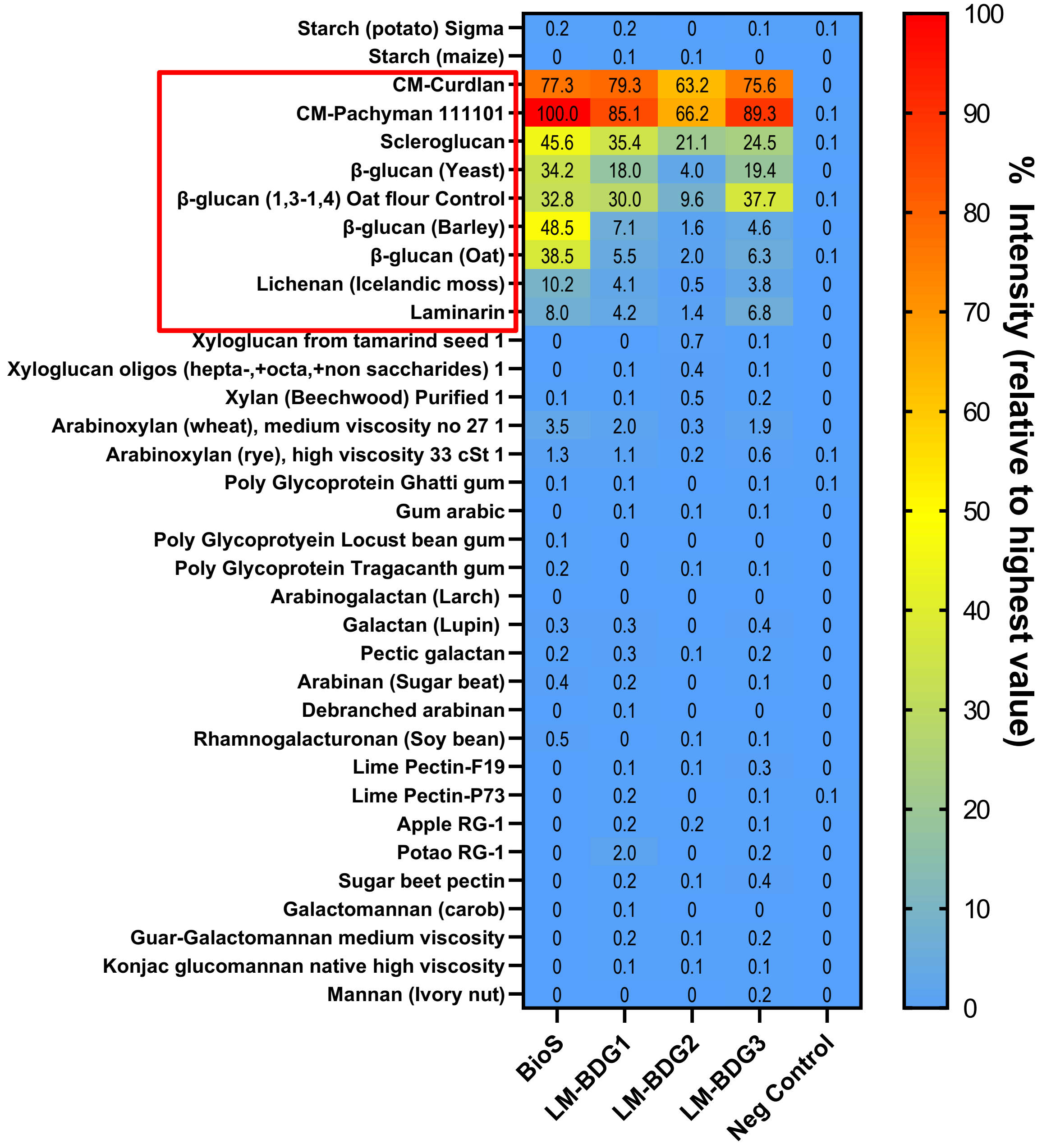


**Fig. S2. Glycomicroarray of callose monoclonal antibodies against polysaccharide standards.** The intensity of the signal relative to the highest value was calculated and presented from low (blue) to high (red) in the table. Commercial substrates with β-(1,3)-glucan linkages are indicated in the red box (including mixed linkage glucans). No substantial binding was observed in components that do not contain β-(1,3)-glucan linkages. Affinity was highest against pachyman and curdlan. BioS has significant binding to mixed linkage glucans which was not observed with LM-BDG2. LM-BDG1 and LM-BDG3 have similar specificity.

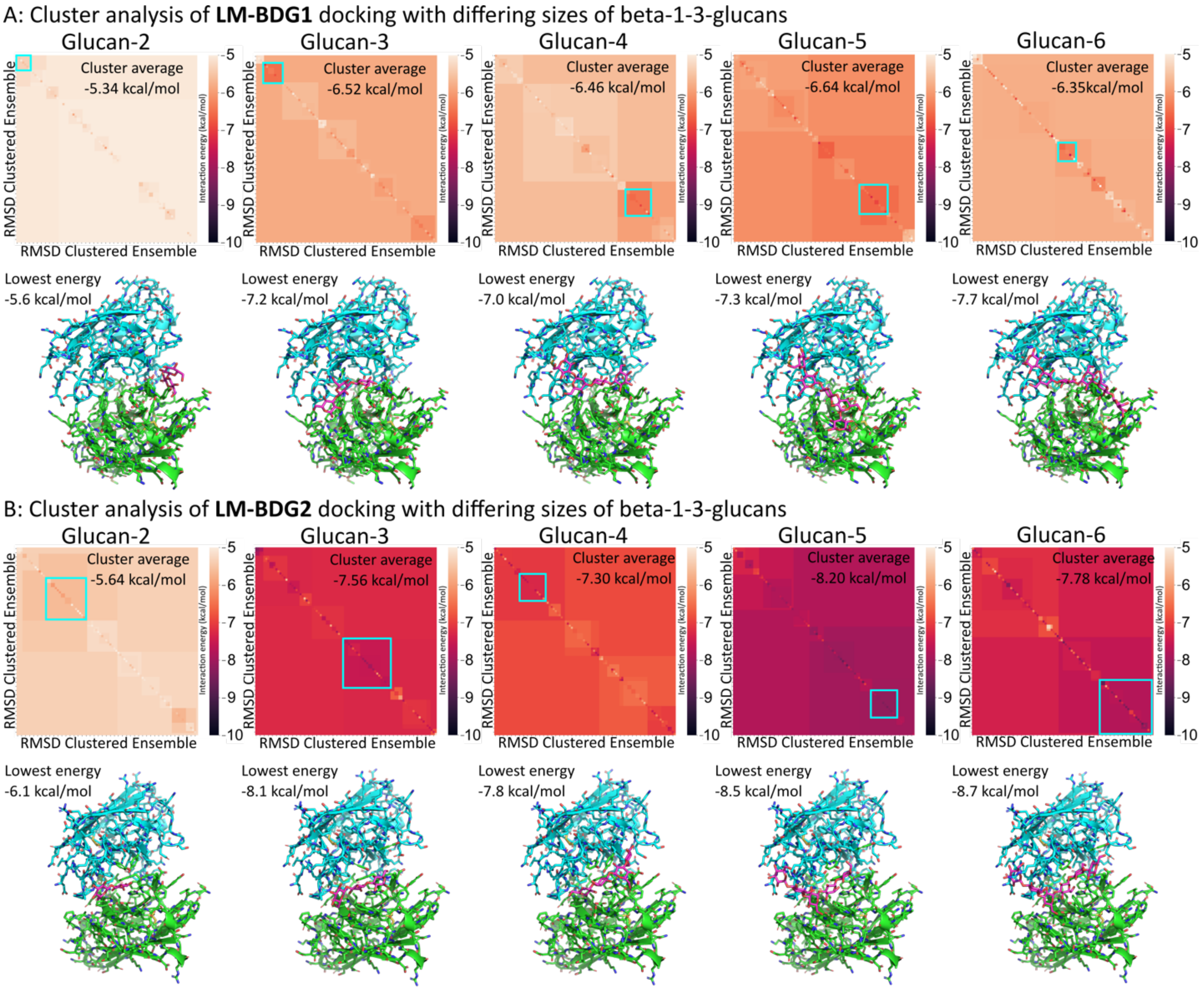


**Fig. S3. RMSD energy landscapes for docked glucans. A**) LM-BDG1 and **B**) LM-BDG2 binding clefts with beta-1-3 glucan oligomers of 2, 3, 4, 5 and 6 residues docked with Vina-Carb. A 2D RMSD based interaction energy landscape was constructed, for which the lowest-energy, high-population energy well was identified (cyan square, cluster average energy displayed, kcal/mol). From this cluster, the lowest energy structure is depicted (antibody in blue and green, glucan in magenta) with interaction energy stated (kcal/mol).

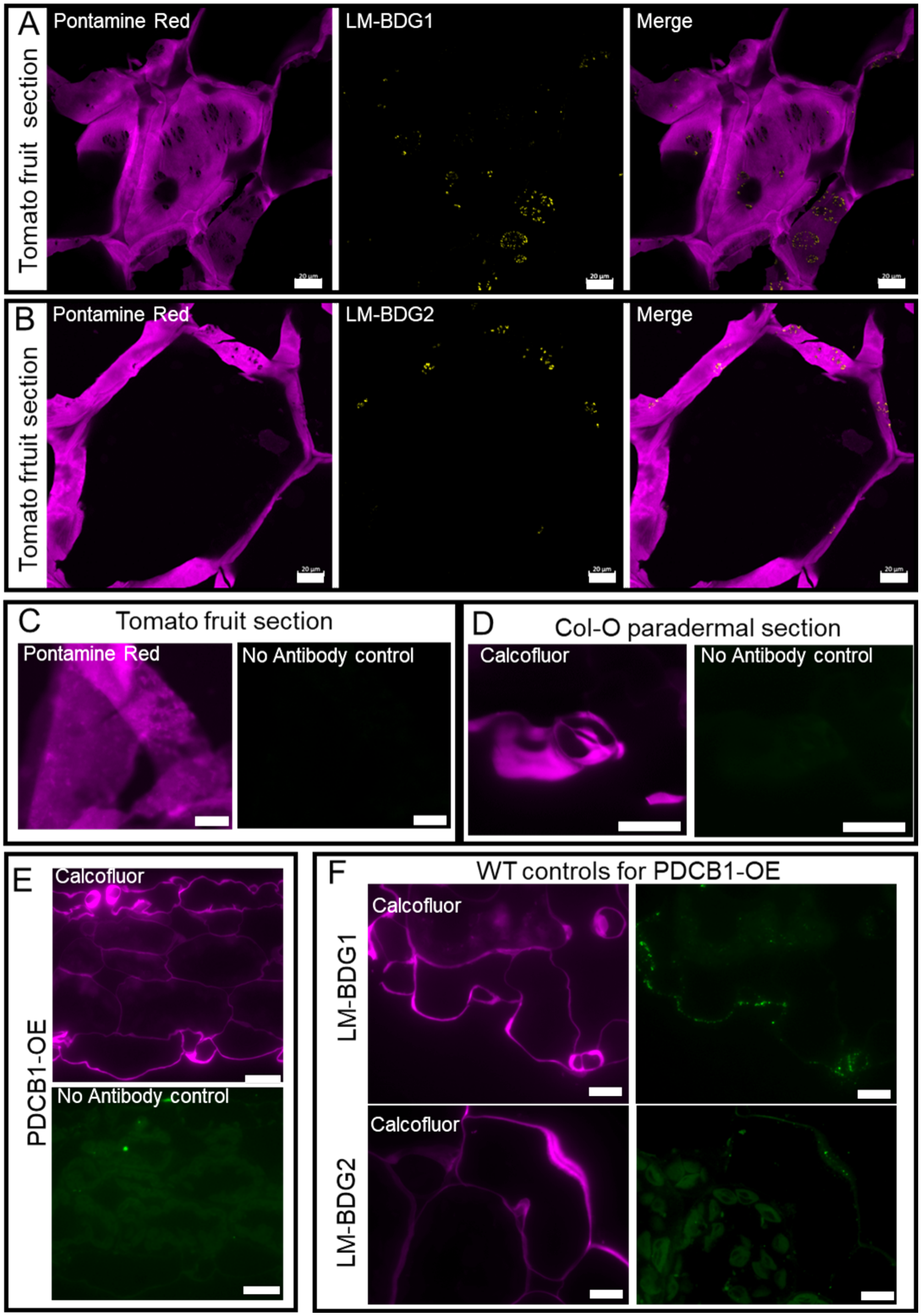


**Fig. S4. Fluorescent immunolabelling of tomato and Arabidopsis sections. (A,B)** Tomato fruit sections were labelled with LM-BDG1 and LM-BDG2 and revealed using Alexa-488-

antirat (green). Pontamine Red (magenta) was used to dye cellulose microfibrils. Note pit fields characterized by low cellulose and high antibody labelling. **(C,D,E)** No-primary antibody controls for tomato (C) Arabidopsis wildtype Col-0 (D) and PDCB1OE, presented in Fig. 2 (E) to capture autofluorescence. **(F)** Wildtype **(**WT) Arabidopsis leaf section collected and labelled alongside PDCB1-OE in Figure 2. LM-BDG1 and LM-BDG2 showed presence of both epitopes but at a lower abundance than in the PDCB1-OE line. Counterstaining of cellulose with calcofluor/pontamine red shown in magenta. Scale bars = 20 µm

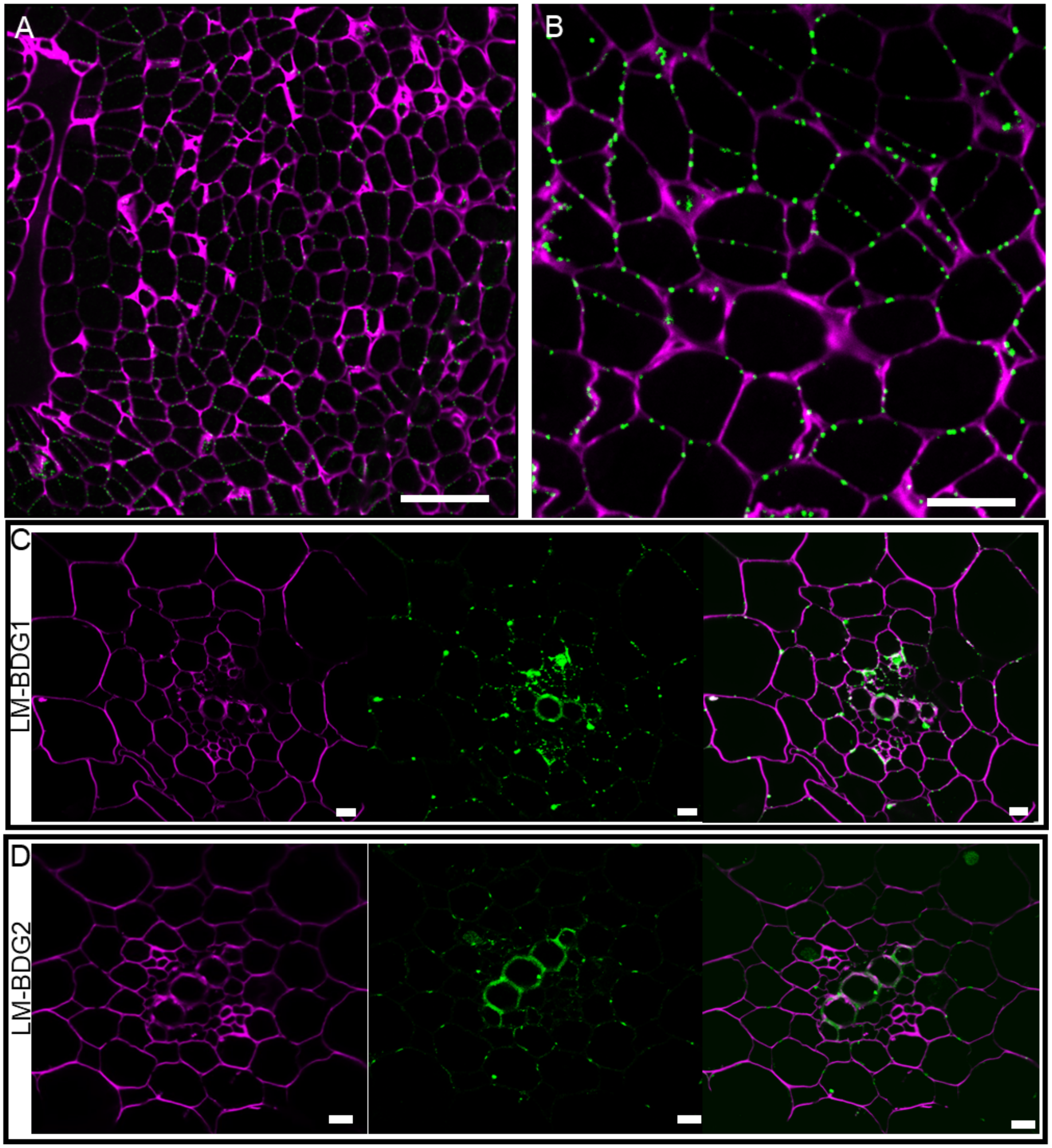


**Fig. S5. Immunofluorescence detection of callose. (A, B)** Immunofluorescence in the shoot apical meristem of Aspen Hybrid T89 using LM-BDG1 and detected with a secondary antibody conjugated with anti-rat-AlexaFluor-555 (green)**. (C, D)** Tobacco root sections were labelled with LM-BDG1 and LM-BDG2 and revealed using anti-rat-Alexa-488 (green). Labelling showed presence of both LM-BDG1 (C) and LM-BDG2 (D). Counterstaining of cellulose with calcofluor white is shown in magenta. Scale bars = 20 µm (A, C, D) and 5 µm (B).

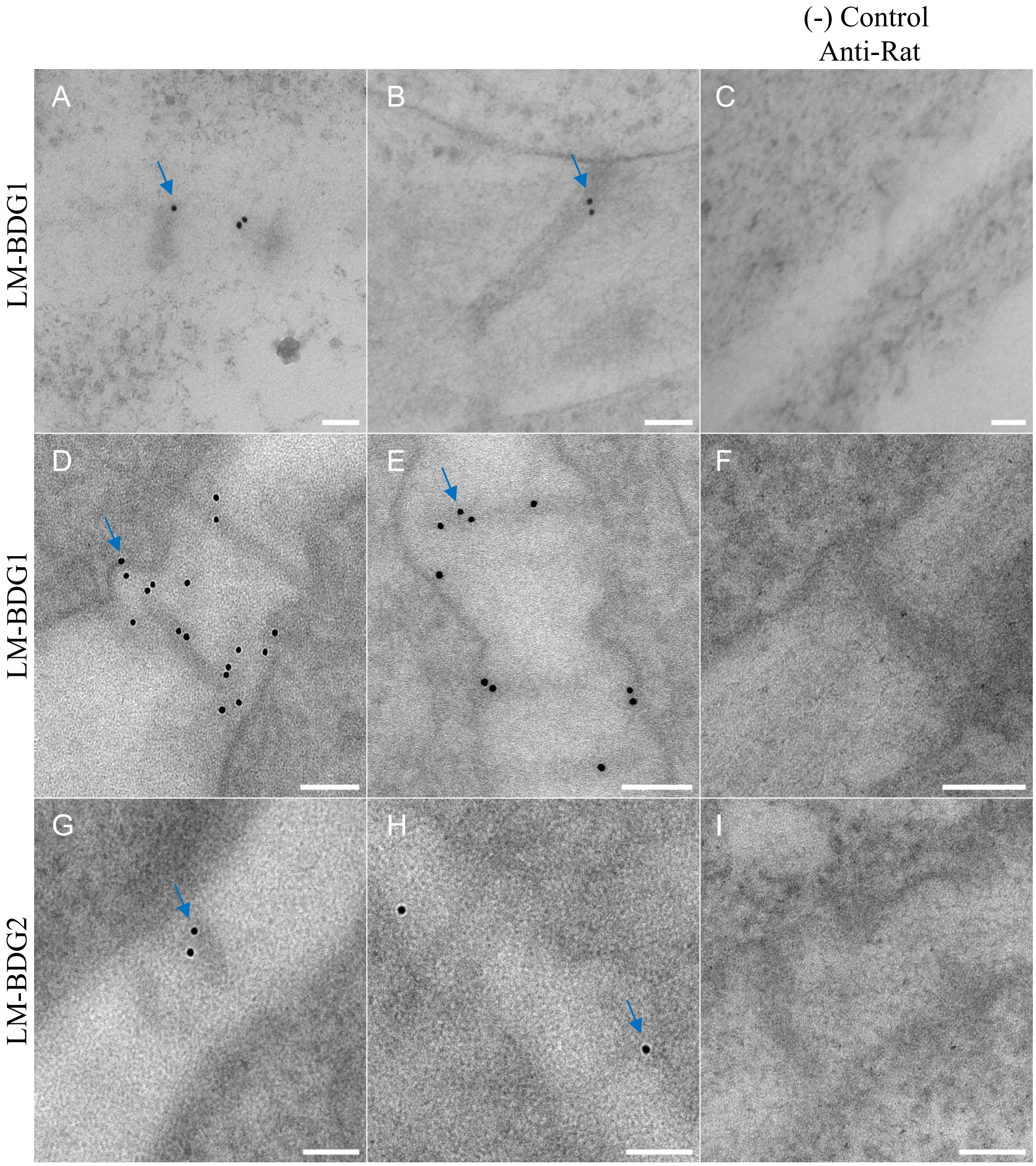


**Fig. S6 Immunogold TEM labelling callose in Arabidopsis cell cultures and aspen buds.** Immunogold EM labelling showed gold-labelled callose (blue arrows, **A-B, D-E, F-G**) detected by LM-BDG1 at plasmodesmata compared to absence of labelling in representative Rat non-immune IgG controls (**C, F, I**). Sections from Arabidopsis culture cells (**A-C**) and the meristem region of Aspen Hybrid T89 (**D-I**) are shown. Scale bar = 100 nm.

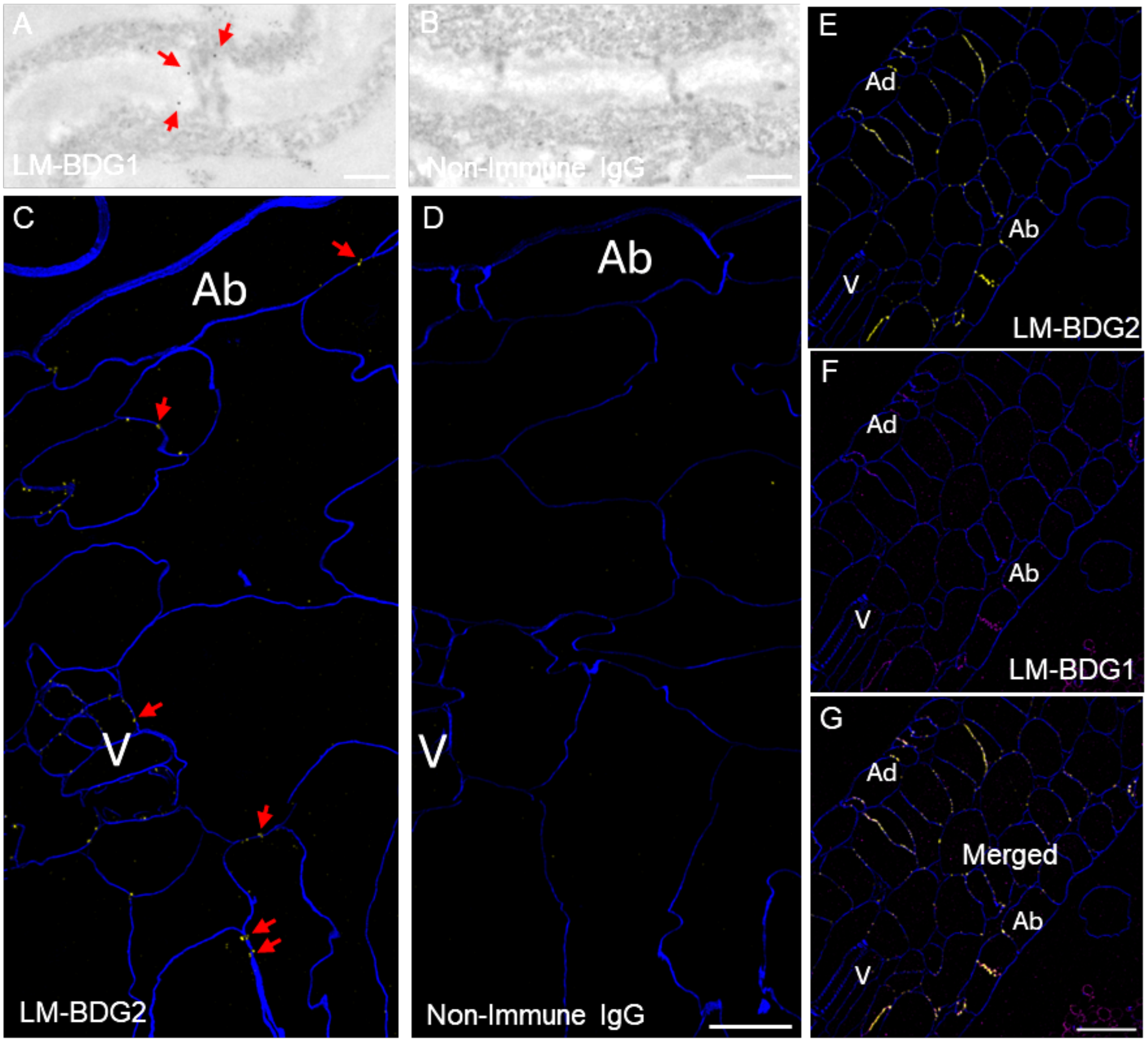


**Fig S7. Immunogold TEM and multiplex immunofluorescence Lattice SIM2 super-resolution microscopy of callose at plasmodesmata (overviews and non-immune controls).** (**A-B**) Immunogold EM labelling in tobacco leaf sections showed gold-labelled callose (red arrows) was detected by LM-BDG1 at plasmodesmata compared to absence of labelling in representative Rat non-immune IgG controls. Scale bar = 200nm. (**C-D**) Tiled large area overview immunofluorescence localization of LM-BDG2 (yellow) at plasmodesmata (red arrows) (**C**) and absence of specific labelling with non-immune Rat IgG negative control (**D**). Calcofluor White cell wall counterstain (blue), adaxial epidermis (Ad), vascular tissue (V). **C** & **D** leaf sections acquired using SIM2 super-resolution microscopy from same samples as Fig. 3 **C-G** and displayed with matched acquisition and brightness contrast settings. Scale bar = 10µm. (**E-G**) Overview Lattice SIM2 super-resolution microscopy immunofluorescence of LM-BDG2 (yellow) and LM-BDG1 (magenta) at plasmodesmata showed some localization overlap and non-overlap adjacency of signals. Calcofluor white cell wall counterstain (blue), adaxial epidermis (Ad), abaxial epidermis (Ab), vascular tissue (V). Scale bar = 20µm.

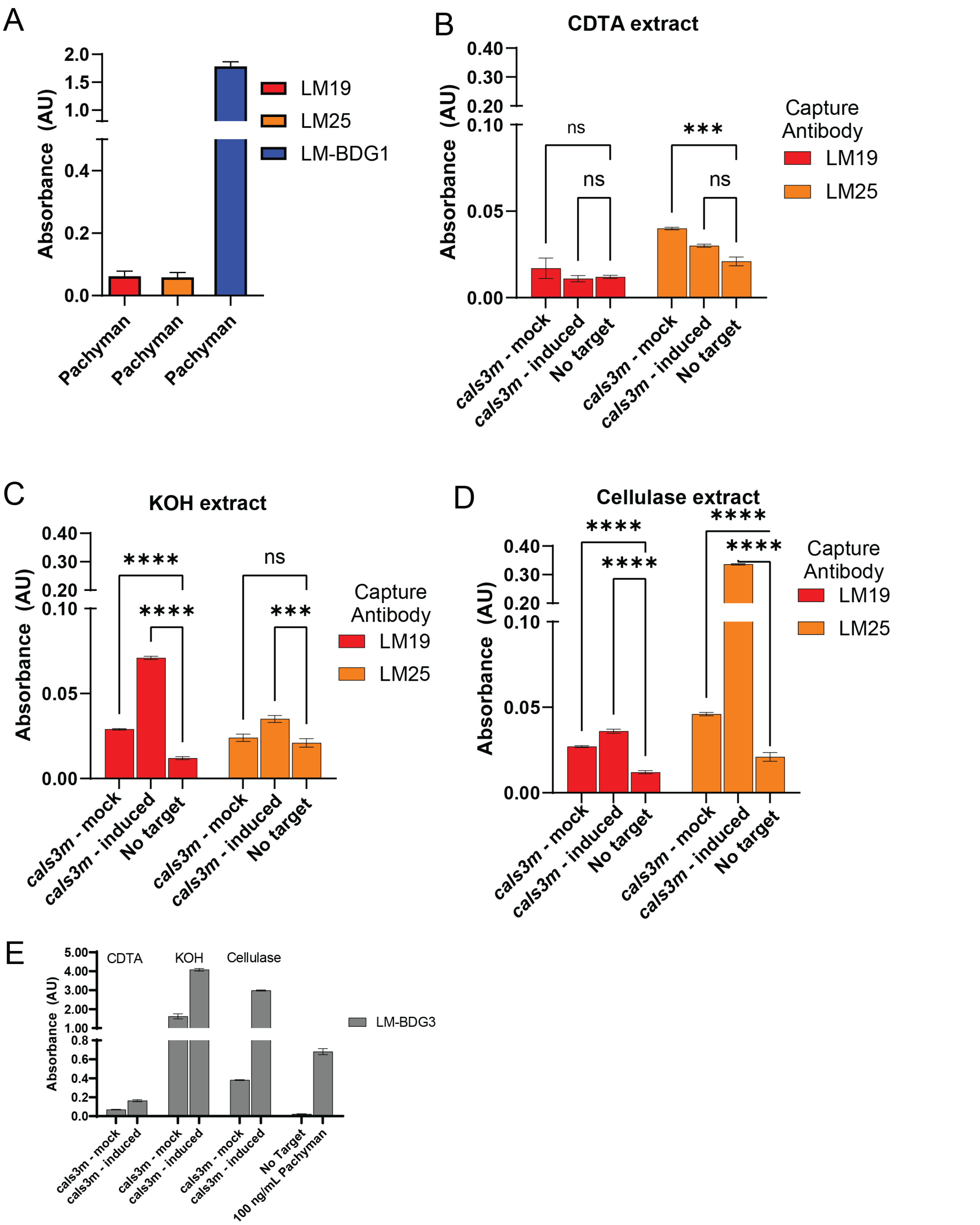


**Fig S8. Sandwich ELISA identifies interactions callose-xyloglucans. (A)** Control experiments carried out using a callose-binding module (CBM43) to capture Pachyman indicate no binding

LM19 (pectin mAb) or LM25 (xyloglucan mAb) but strong signal when using LM-BDG1. Error bars represent standard deviation from n=4 pools of 4 plants each. **(B-D)** Sandwich ELISA were also carried out in reverse: using LM25 and LM19 to capture respective antigens and a purified HRP-tagged version of LM-BDG1 to detect calloses that co-precipitate with pectin and xyloglucans in the different cell wall extracts (CDTA, KOH and cellulase digested fractions). Callose co-precipitation was higher (higher Absorbance) in estradiol-induced *cals3m* (i*cals3m*) in relation to mock (DMSO) un-induced control. No target (no cell wall extract) blank controls are also represented. Note callose is detected when precipitating pectin in the KOH extract (**C**) and mainly with xyloglucans in the cellulase extract (**D)** suggesting interactions between these polymers. (**E**) To confirm callose immuno-detection by LM-BDG1-HRP, sandwich ELISA were also carried out using LM-BDG3 as a capture antibody. Increases in calloses by i*cals3m* were observed in all extracts. Pachyman was used as positive control. Error bars show standard error (SEM) from n=3. Statistical significance: *** indicates $p<0.005$ *** indicates $p<0.0005$ determined by 2-way ANOVA.
